# *Avant-garde*: An automated data-driven DIA data curation tool

**DOI:** 10.1101/565523

**Authors:** Alvaro Sebastian Vaca Jacome, Ryan Peckner, Nicholas Shulman, Karsten Krug, Katherine C. DeRuff, Adam Officer, Brendan MacLean, Michael J. MacCoss, Steven A. Carr, Jacob D. Jaffe

**Affiliations:** Broad Institute of MIT and Harvard, Cambridge; University of Washington Genome Sciences, Seattle, WA

## Abstract

Data-Independent Acquisition (DIA) is a technique that promises to comprehensively detect and quantify all peptides above an instrument’s limit of detection. Several software tools to analyze DIA data have been developed in recent years. However, several challenges still remain, like confidently identifying peptides, defining integration boundaries, dealing with interference for selected transitions, and scoring and filtering of peptide signals in order to control false discovery rates. In practice, a visual inspection of the signals is still required, which is impractical with large datasets. *Avant-garde* is a new tool to refine DIA (and PRM) by removing interfered transitions, adjusting integration boundaries and scoring peaks to control the FDR. Unlike other tools where MS runs are scored independently from each other, *Avant-garde* uses a novel data-driven scoring strategy. DIA signals are refined by learning from the data itself, using all measurements in all samples together to achieve the best optimization. We evaluated the performances of *Avant-garde* with a calibrated sample using spiked-in standards in a complex background, a phospho-enriched dataset (Abelin *et al*, 2016), and two complex hybrid proteome samples for benchmarking DIA software tools. The results clearly showed that *Avant-garde* is capable of improving the selectivity, accuracy, and reproducibility of the quantification results in very complex biological matrices. We have further shown that it can evaluate the suitability of a peak to be used for quantification reaching the same levels of selectivity, accuracy, and reproducibility obtained with manual validation.

## Introduction

Quantitative proteomics is a fundamental tool to decode the complexity and the dynamics of the proteome. Data-Independent Acquisition (DIA) is a family of newly-developed LCMS acquisition methods combining the unbiasedness of Data-Dependent Acquisition (DDA) with the reproducibility, sensitivity, and accuracy of targeted methods^1–4^. In DIA, MS instrumentation co-isolates and fragments multiple peptides, either in sequential isolation windows traversing an *m/z* range, or all at once^1–9^. DIA has the potential to comprehensively analyze all peptides in a sample that are above the instrument’s limit of detection.

DIA data are quantified with a chromatogram-based approach. For each peptide, several transitions (precursor/fragment ion pairs) are monitored over time, producing a set of chromatographic peak traces. Peak area is integrated and used as a proxy for analyte abundance. Ideally, all transitions of a given analyte should have: 1) the same elution peak shape, 2) relative areas mirroring the relative intensities found in their reference spectrum from a library, 3) a low mass error, and 4) consistency across all MS runs being compared. However, due to the complexity of DIA data, it is difficult to obtain signals that correspond to this archetype, and data analysis remains challenging.

Several tools have been developed to analyze DIA data^10^. Each one can produce a different set of detectable peptides and quantitative results, even with standardized samples and data sets^11^. This variability is introduced by differences at all stages of data analysis (i.e. raw data processing, protein database search, peak detection, transition selection, chromatogram extraction, peak integration, and statistical analysis), each of which can affect detection and quantification.

Most tools focus on statistical validation of peptide detection (using target/decoy approaches^12, 13^) but do not address the *quantitative suitability* of the signals extracted. Targeted analyses of DIA data begin with spectral libraries, which may be built from a single, fractionated “master sample” using narrow isolation window DDA methodology. These practices mask the complexity found in real DIA data and do not anticipate interferences present in real biological samples, especially when perturbations are introduced. Therefore, transitions selected from spectral libraries may not be suitable for quantification in actual biological sample sets. In practice, further curation of signals is required for rigorous quantitation. During curation, an expert visually inspects the data, removes transitions subject to interference, and manually corrects peak integration boundaries. However, time-consuming manual curation is impractical with large datasets, and produces subjective user-dependent results. Curation is thus often omitted and the output of these tools is used at face-value for downstream analysis.

Missing values are also a problem in DIA approaches. They can have a biological origin (e.g. peptides truly not present), a technical origin (e.g., peptide loss during sample processing), or a computational origin, (e.g., failure to assign the correct signal to the respective peptide). The latter reason can be due to retention time prediction or alignment models that improperly impute chromatographic boundaries of the analytes in real samples. Missing data can also originate from non-curated data. A peptide subject to interference might fail to be identified in a subset of samples (e.g. indistinguishable from a decoy peptide), thus creating missing values even though the peptide is present. However, if another set of interference-free transitions had been used, this peptide might be detected in all samples, subsequently providing accurate and reproducible quantification.

The issues discussed above motivated us to create a tool for automated targeted MS data curation. Here we present *Avant-garde* (AvG), a modular tool meant to polish the results of DIA and PRM analysis tools. Building upon earlier work on DIA and PRM data optimization^14^, AvG refines DIA signals to reach the highest possible levels of sensitivity, selectivity, and accuracy. AvG refines peak detection, adjusts peak boundaries, removes transitions subject to interference, eliminates noise, and estimates the FDR of analytes for *quantitative suitability*. Unlike other tools where MS runs are scored independently from each other, AvG uses a novel ensemble data-driven scoring strategy. DIA signals are refined by learning from the data itself, using all measurements in all samples together to achieve the best optimization.

## Results

### Principle of Avant-garde

AvG is a tool designed for automated data curation, meant to complement common DIA analysis tools such as mProphet^12^, OpenSWATH^13^, DIA-Umpire^15^, EncyclopeDIA^16^, and Specter^17^. To ease use and adoption of AvG, we have chosen Skyline^18^ to extract chromatogram data as a vendor-independent and user-friendly tool. It enables data visualization and provides a common framework to refine results from different upstream tools. Skyline requires only the peptide sequences and peak integration boundaries determined by these tools.

AvG uses the chromatogram data and employs three independent modules to refine the data (Fig. 1). First, a transition refinement module improves the choice of transitions to eliminate interferences and reduce noise. Second, a peak refinement module adjusts integration boundaries without the need for spiked-in retention time peptides. A third module scores peaks using a number of intuitive metrics and estimates the false discovery rate (FDR) for quantitative suitability. The refinement results and scoring metrics are then imported back into Skyline.

**Figure 1:**
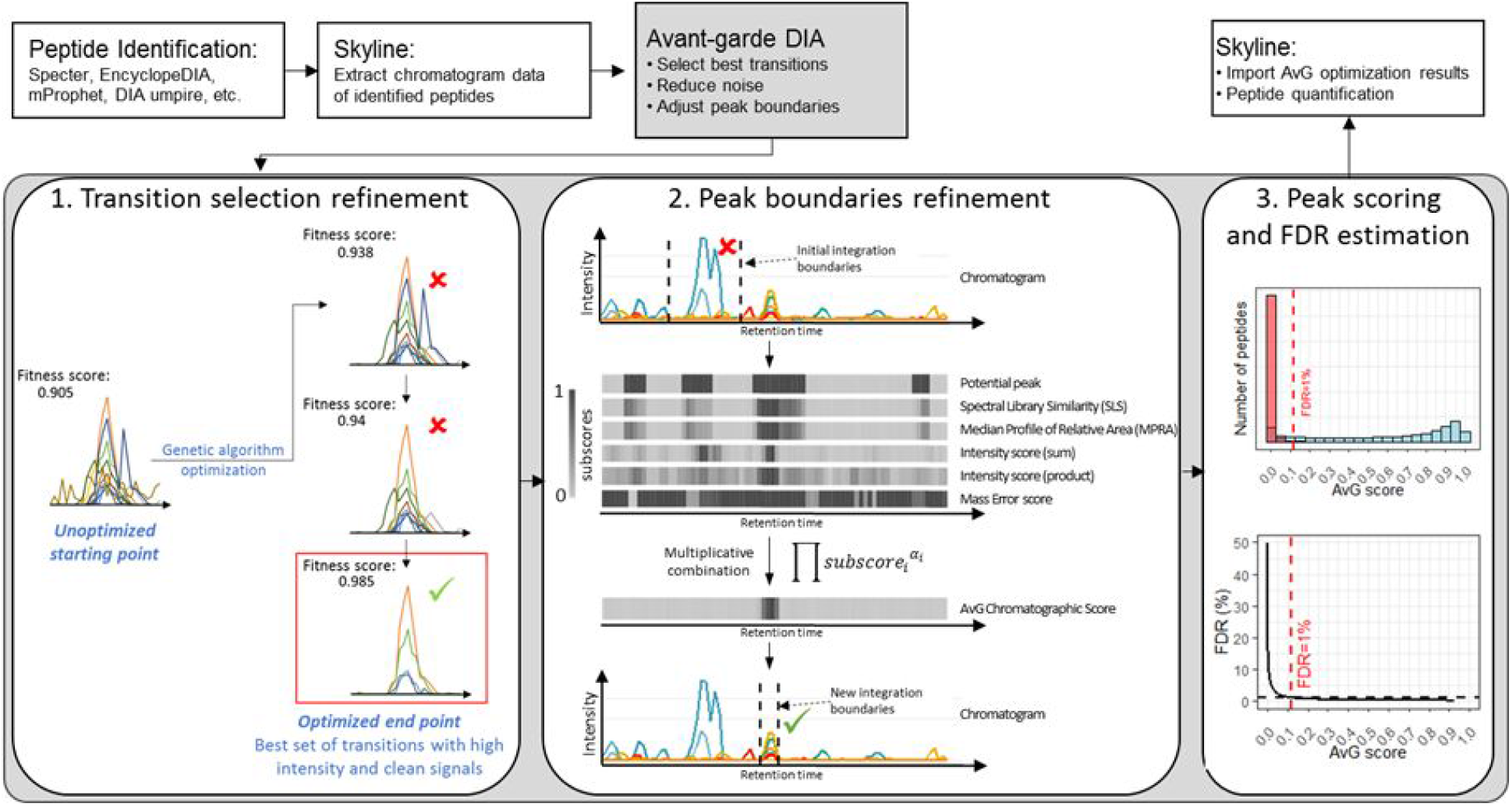
*Avant-garde’s role* in data analysis and modular scheme. *Top:* AvG is employed downstream of independent DIA identification engines, and depends on Skyline to extract chromatogram data of detected peptides. The output of AvG (suitable transitions, chromatographic boundaries, and scoring metrics) is reimported to Skyline to produce curated quantitative data. *Bottom:* AvG is composed of three modules. Module 1 curates transitions to reduce noise and remove interference using a genetic algorithm, assigning a final quality metric to the selected set (*AvG fitness score*). Module 2 refines peak integration boundaries. AvG calculates chromatographic subscores at each time point in the raw data, and combines them as a weighted product (*AvG chromatographic score*). The maximum value of this score corresponds to the most likely retention time of the analyte. Module 3 scores peaks (*AvG score*), filters the data and estimates the FDR for quantitative suitability.

Like other tools, AvG assigns and quantifies peptides using composite scores (built from subscores) as the basis for quality filtering of results and estimation of the FDR. Each module of AvG produces its own composite score: the “*AvG fitness score*” for transition selection, the “*AvG chromatographic score*” for peak integration boundaries, and the “*AvG score*” for the final scoring of peaks and FDR estimation. However, its composite scores are calculated as the product rather than the sum of its subscores. This approach avoids allowing any *single* subscore to push the composite score over an arbitrary “significance” threshold employed to control the FDR. Uniquely, AvG calculates its module scores in an ensemble-driven manner, curating transitions and peak boundaries while *considering data from all samples in a set*.

The scoring strategy is designed to produce very conservative results. AvG penalizes peptides with any single metric that indicates poor quality. This scoring mechanism imposes strong penalties on transitions subject to interference. A high final AvG score ensures that minimal interference is present and that the signals are suitable for quantification.

### Automated refinement of transition selection by a genetic algorithm

The transition refinement module is based on a genetic algorithm (Fig. 1), which is a machine learning method designed for solving optimization problems. Its operating principle mimics natural selection and biological evolution, and efficiently samples a large number of combinations without being exhaustive. To select the best set of transitions for each peptide, we start by extracting a large number of transitions for each (*<u>at least</u>* 5-10 per peptide). These transitions may be subject to interference (Fig. S1). For each step (or generation), the genetic algorithm selects several random subsets of transitions. The algorithm then scores each one with a fitness function and selects the best-scoring solution(s) as the starting population for the next generation. Over successive steps, the population “evolves” towards an optimal solution, as the score of the fitness function increases until it reaches a stable maximum. The corresponding solution is the most suitable set of transitions for quantification.

The genetic algorithm maximizes the AvG fitness score (Supp. Methods and Fig. S2). The subscores fall into two categories: run-specific and dataset-wide scores. There are three run-specific subscores: 1) the mass error score penalizes differences between observed and expected masses of transitions, 2) the Spectral Library Similarity (SLS) score compares the relative intensities of transitions to a reference spectrum using a transformation of the dot product, and 3) the Peak Shape Similarity (PSS) score evaluates the correlation of elution profiles of transitions. The PSS score is extremely sensitive for detection of interference.

The median profile of relative areas (MPRA) subscore is a dataset-wide score (Supp. Methods). In a nutshell, the MPRA measures how similar the signals in a given run are to all other runs in the entire dataset. It evaluates the similarity of the relative peak areas of transitions in one run to the median profile calculated on all the runs in the dataset.

For each run, the composite score is a combination of run-specific and dataset-wide scores. This ensures that the final solution is the best possible compromise considering all runs in the dataset and is not influenced only by a small number of high-scoring runs. It also allows the identification of problematic runs. Finally, the *AvG fitness score* is calculated for each peptide in the dataset by calculating the mean of all combined scores for each run. Because the *AvG fitness score* is ensemble-driven, the genetic algorithm evolves towards noise-free signals, reduced interference, and consensus in quantifiable transitions across the entire dataset.

### Automated refinement of peak integration boundaries

AvG uses the curated transitions to adjust peak boundaries without the need for retention time alignment. Several chromatographic subscores are calculated at each time point in the raw data (Fig. 1). The *AvG chromatogram score* is also calculated in an ensemble-driven manner as a product of subscores (Supp. Methods). The objective is to penalize peptides with any single metric that indicates poor quality. The maximum value of the AvG chromatogram score corresponds to a peak’s most likely retention time.

Fig. 1 (box 2) shows an example of the cumulative effect of transition curation and peak boundary refinement for a peptide. Initially, the integration boundaries (from Skyline) did not correspond to the real chromatographic peak. AvG focuses on signals that are potential peptide peaks, where for each point at least 3 non-zero transitions are observed. For potential peaks, the SLS and the MPRA are calculated. In the example, some subscores (SLS, MPRA, and the mass error) are low in the time range corresponding to the initial peak integration boundaries. The *AvG chromatographic score* has a low value close to zero in that time window, even though the intensity scores are high. However, the maximum value of the *AvG chromatographic score* corresponds to the real retention time of the analyte. The peak boundaries are then defined around this time, guided by the *AvG chromatographic score* and the *Intensity (sum) subscore*. Prior knowledge of approximate retention time can improve the speed of this module, but is not a requirement (see Supp. Fig. 3 for the same example over a wider time window). When a peptide is not present in a sample for biological or technical reasons, AvG still integrates the signal around the maximum value of the ensemble-driven *AvG chromatographic score*.

### AvG identifies DIA signals with high quantitative suitability

After curation of transitions and peak boundaries, all signals are scored again with the third module to assess overall quality, and suitability for quantification (Supp. Methods). As described above, the scoring mechanism was designed to discriminate poor-from high-quality signals with respect to quantitative suitability. To evaluate the selectivity, precision, accuracy, and FDR of AvG results, a five-point calibration curve (5 samples analyzed in triplicate) was built using a background of HEK293T whole cell digest spiked with 95 synthetic phosphopeptides. The calibration curve ranged from 6.75 ng to 108 ng of total synthetic peptide mixture spiked into 1 μg of HEK293T digest per injection. Each sample was measured in triplicate on a Q-Exactive HF using a DIA method (Supp. Methods).

First, we determined the FDR for *quantitative suitability* after curation with AvG using a target-decoy approach. To do this, we randomly selected 1000 human peptides that were identified in previous DDA runs of the same digest. Their signals and their corresponding shuffled-sequence decoys were extracted from DIA runs and were automatically curated using AvG.

The distributions of each subscore (Fig. S4) discriminate between targets and decoys with varying performance. However, a weighted product of the subscores produces an extremely good separation of the distributions. We empirically determined the exponent weights for the multiplicative combination of the AvG subscores. However, we verified that the results obtained with our weights matched the results with weights obtained by linear discriminant analysis (Supp. Methods and Fig. S5) with respect to overall sensitivity. Most decoys have an AvG score close to 0 (98% < 0.05) and the cutoff to obtain an FDR below 1% was a value equal to 0.11.

AvG additionally enforces minimum thresholds for each subscore (SLS > 0.7, mass error score > 0.7, PSS > 0.85, MPRA > 0.9) to remove poor signals (Fig. S4). Filtering by these thresholds does not alter the total number of analytes marked as suitable for quantification and the FDR remained below 1% (Fig. S6). A final *AvG score* threshold of 0.1 was adopted. FDRs computed using these thresholds were always < 1.0%.

### AvG ensures high accuracy and precision

The precision and accuracy of AvG were evaluated using the dilution series of the 95 synthetic phosphopeptides that were spiked into the HEK293T digest. A typical AvG performance on the spiked-in synthetic peptide S[+80]LTAHSLLPLAEK is shown in Fig. 2A. Skyline’s initial extraction of signals from this peptide is incorrect in some runs, due to misassignment of peak boundaries. These aberrant signals cause departure from the expected linear relationship between concentration and peak area (*r*^2^ = 0.44, Fig. 2A, top). After curation by AvG, the expected linear relationship is recovered (*r*^2^ = 0.99, Fig. 2A, bottom).

**Figure 2:**
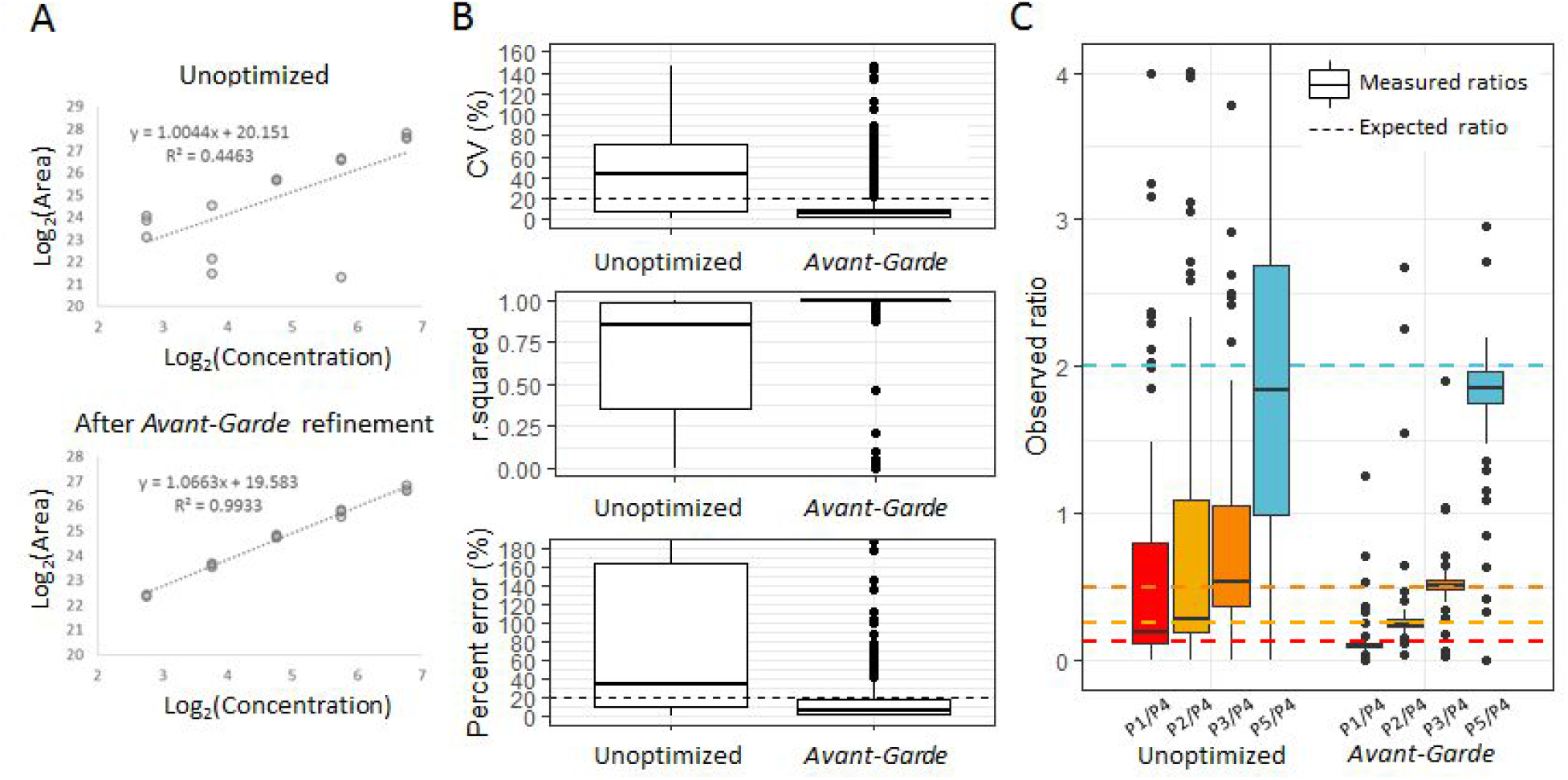
AvG improves quantitative figures-of-merit in a calibration curve. 95 synthetic phosphopeptides were spiked into a HEK293T whole cell digest to create a 5-point calibration curve (5 samples were analyzed in triplicate). (A) Calibration curve before (top) and after (bottom) curation by AvG for peptide S[+80]LTAHSLLPLAEK. In this case, AvG automatically corrected the peak boundaries for this peptide improving curve linearity. (B) Figures-of-merit summarising the results for all synthetic peptides, pre- and post-optimization: % CV of triplicates, *r*^2^ values of linear fits, and absolute percent error of measurements relative to the known concentration. Dashed lines indicate 20% thresholds. The box plot elements are: center line, median; box limits, upper and lower quartiles; whiskers, 1.5x interquartile range; points, outliers (C) Expected vs. observed ratiometric quantification, pre- and post-optimization. P1 to P5 represent the points of the calibration curve in increasing order of concentration. The ratios between the mean area of each calibration point to the mean area of the fourth calibration point (P4) are shown here. The dashed lines represent the expected ratios (0.125, 0.25, 0.5 and 2) and the boxplots show the distribution of the measured ratios. The boxplot elements are the same as described for panel B.

There was marked improvement for aggregate measurements of all 95 synthetic peptides after application of AvG (as compared to initial, unoptimized Skyline extractions) in several figures of merit (Fig. 2B). The precision, measured by the CVs of triplicates, improved from 43.2% to 5.6%. The correlation coefficient between peptide concentration and peak area improved from *r*^2^=0.85 to *r*^2^ =0.99. Improvements in the fraction of measurements with less than 20% absolute error (our definition of accuracy, see Supp. Methods) were also evident. Finally, the relative quantification accuracy was also evaluated. We calculated the ratios between the mean area of each calibration point to the mean area of the fourth calibration point (P4 in Fig. 2C). The distribution of measured ratios after AvG refinement is clearly much tighter and closer to the expected values.

### AvG results are concordant with expert manual curation

To further evaluate the performance of the automated data curation, we applied AvG to a reduced-representation phosphoproteomics dataset obtained for our LINCS project^19^. This dataset, acquired in DIA mode, had previously been manually curated by an expert in our laboratory, which we consider the gold-standard against which other approaches were compared. We curated data across 96 samples for 95 phosphopeptides for which isotopically-labeled heavy peptide counterparts were present. For the “unoptimized” analysis, the 5 most intense transitions from the spectral library were chosen and the peak boundaries were defined by Skyline. For the optimized version, all possible b- and y-ions above b_4_ and y_4_ were extracted and subjected to further curation by AvG. AvG was run in two modes: 1) “open” curation, where no subscore or composite score filters were applied, and 2) “filtered” curation, where subscore filters were introduced (see discussion of filtering above).

The comparison of light-to-heavy ratios between the manually curated and unoptimized analyses (Fig. 3A) had many points deviating from the ideal x=y line. After “open” curation by AvG (Fig. 3B), many fewer points deviated from this line. The disagreements that remained could be explained by peptides where AvG chose different transitions than the manual curator, producing discrepant light-to-heavy ratios. These differences were enhanced if either the light or the heavy peptide had low intensity. In that case, any small change in the signal would have a large impact on the ratio. The results of filtered curation correlated even better with manual curation (r=0.99, Fig. 3C). Signals creating discrepancies in the open curation analysis were filtered out showing that they were derived from low-quality data.

**Figure 3:**
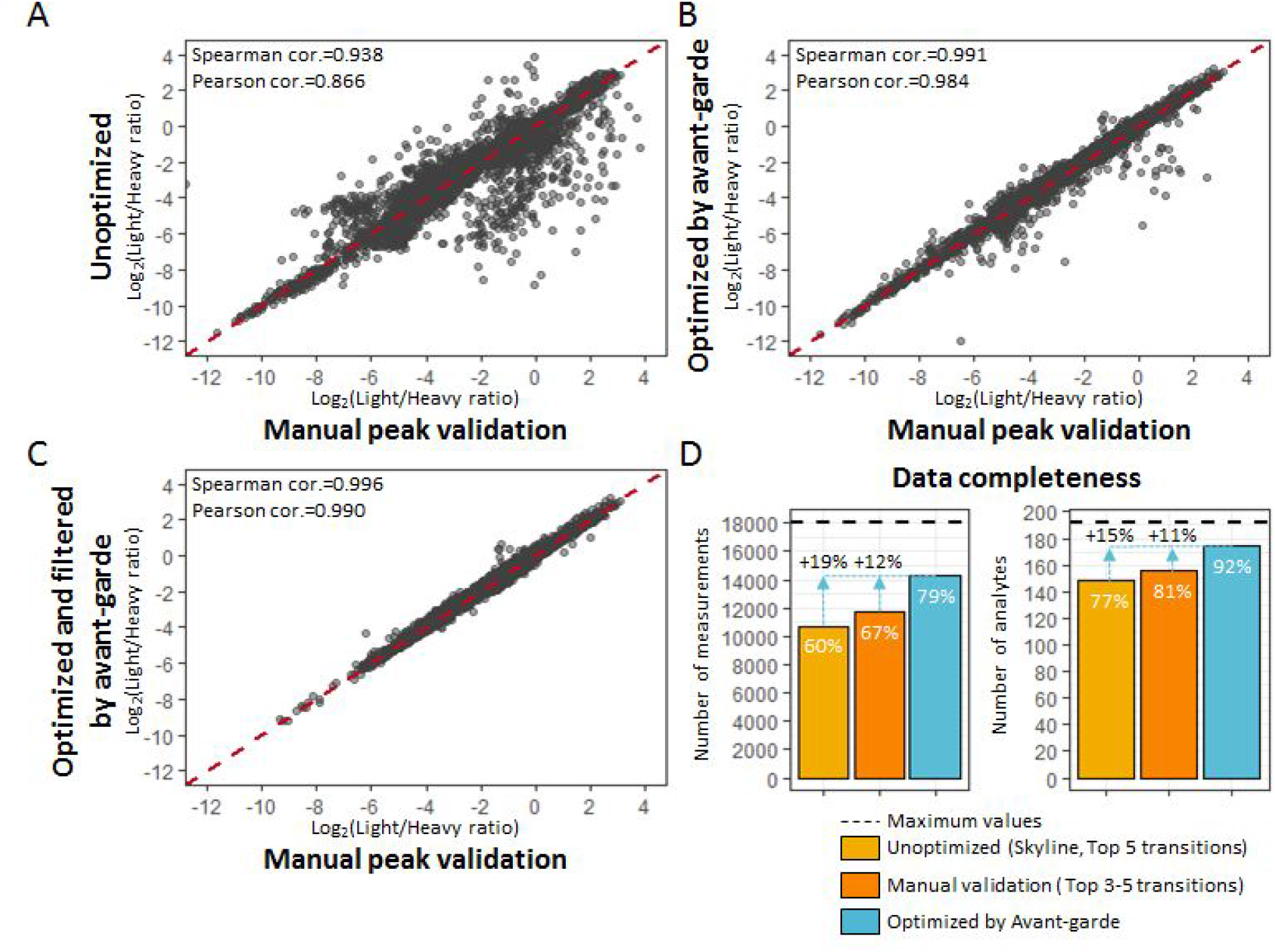
Avant-garde equals the performances obtained by expert visual inspection and manual validation. We focused on 95 phosphopeptides, and their isotopically labeled heavy peptide counterparts, analyzed in a cohort of 96 phospho-enriched samples. The dataset was initially analyzed using Skyline and manually curated by an expert. The scatter plots compare results of light-to-heavy ratios of the (A) unoptimized dataset, (B) the AvG “open” curation dataset, and (C) the AvG filtered curation dataset to the manually curated dataset. (D) Data completeness measured after filtering the data for quantitative suitability at the measurement and at the analyte level.

Additionally, AvG improved the data completeness. Analysis of 190 precursors and 18240 individual measurements was theoretically possible (95 peptides x 2 isotopic label states x 96 samples). AvG improved the data completeness over unoptimized analysis and even manual curation (19% and 12%, respectively, at the measurement level, Fig. 3D). Overall, the data curation by AvG enabled the quantification of 92% of all peptides with data completeness of 79%. AvG performed its curation of this dataset in < 1 hr of unsupervised time, while typically it takes a manual curator > 10 hours of “hands-on” work.

### Evaluation of AvG with LFQBench

We asked whether AvG could further improve quantification when applied to the leading DIA benchmarking dataset in the field, LFQBench^11^, as compared to the many tools with which it has already been analyzed. This dataset was collected on a time-of-flight mass spectrometer, resulting in different data characteristics (resolution, mass accuracy, baseline noise level) than the Orbitrap-class data on which AvG was developed.

The example shown in Fig. 4 compares the basic Skyline analysis to the AvG curation of the LFQBench HYE110 dataset, acquired with a SWATH method with 64 variable m/z windows^11^. The two samples are each mixtures of three complex proteomes - *E. coli*, human and yeast - formulated as shown in Fig. 4A. Three expected ratios are possible when comparing sample A to B (0.1, 1 and 10 for the *E. coli*, human and yeast peptides respectively). The results extracted using Skyline show that a large number of ratio data points deviate from the expected values (Fig. 4B). Even the *centers* of the distributions of ratios for the non-human proteomes did not match their expected values. The median percent errors were 373% for *E. coli* and 84% for yeast. These deviations also affected the precision of the measurement, calculated for triplicate values (median CV of 15.1%, 7.3%, and 19.3% for *E. coli*, human and yeast peptides respectively, Fig. 4D).

**Figure 4:**
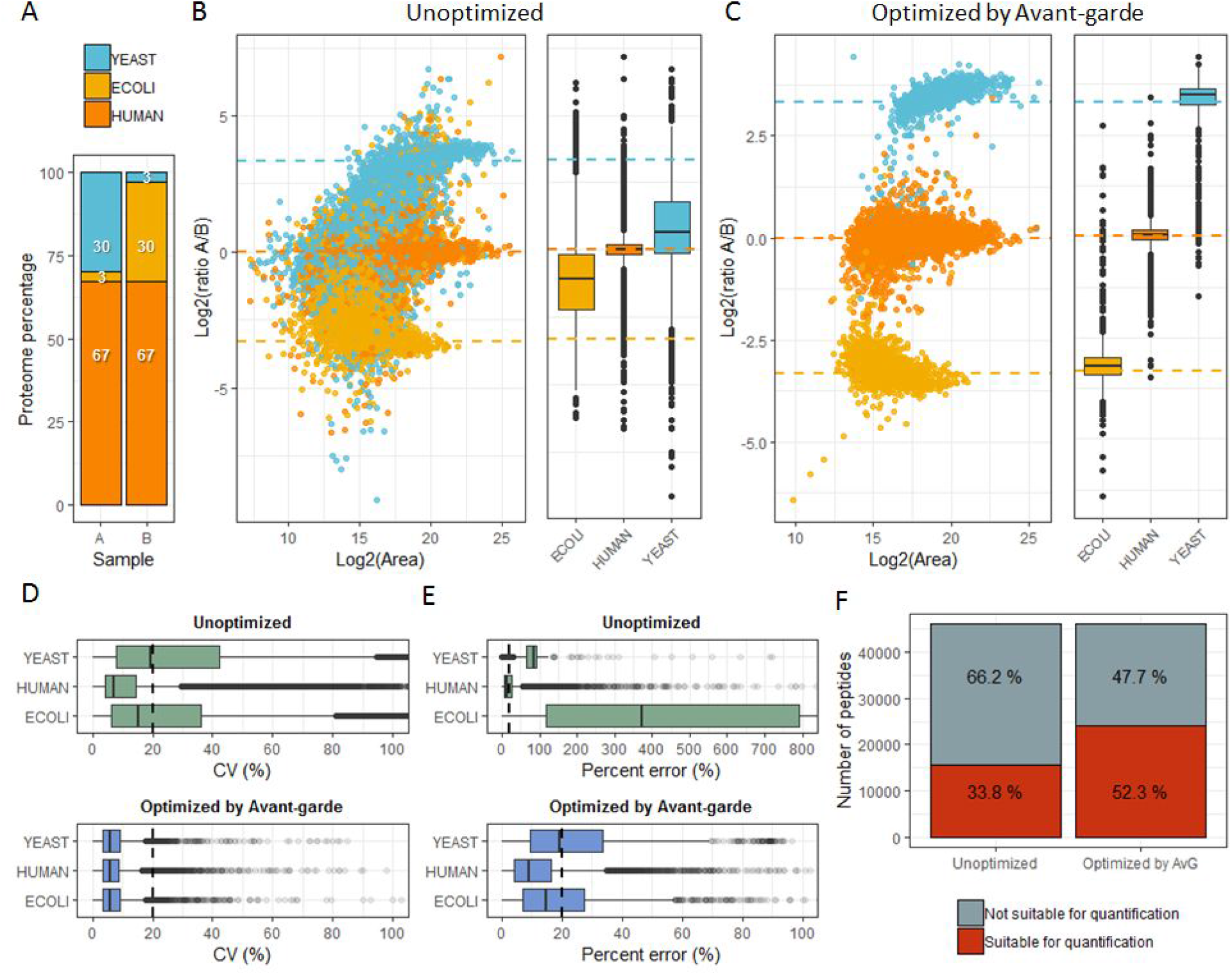
Evaluation of AvG with LFQBench data. (A) The composition of the LFQBench samples by species proteome. (B, C) Results of the relative quantification and distribution of the experimental ratios obtained in the unoptimized dataset (B) and the AvG-curated dataset (C). Each dot represents a ratio calculated for a given peptide in a given run. The dashed lines represent the expected ratios. (D) The coefficient of variation and (E) the percent error for each proteome for the uncurated (top) and curated (bottom) dataset are shown. The vertical dashed lines demarcate the 20% threshold. The box plot elements are: center line, median; box limits, upper and lower quartiles; whiskers, 1.5x interquartile range; points, outliers. (F) The percentage of measurements meeting the criteria for quantitative suitability (AvG module 3) is shown before and after curation with AvG.

After curating and filtering the data with AvG, the ratio distributions were much closer to the expected values (Fig. 4C). The median A/B ratio for each proteome was 0.11 ± 0.015 for *E. coli*, 1.03 ± 0.09 for human, and 11.24 ± 1.91 for yeast. The precision improved dramatically (median CV of 5.8%, 5.7 %, and 6%; Fig. 4D), as did the accuracy (median % error 5% and 19% for *E. coli* and yeast respectively; Fig. 4E).

AvG produced very conservative results. The total number of reported peptides after curation was lower than initially reported by the upstream tools. We wanted to evaluate the quality of the data in both pre- and post-curation analyses. To do this, we “marked” peptides as suitable for quantification by independently applying AvG’s third scoring/filtering module (Supp. Meth., and Figs. S7 and S8). In the unoptimized analysis, only 34% of over 240,000 initially reported measurements were marked as suitable according to our metrics. After the signal curation by AvG this fraction increased to 52%. This implies that AvG improves the number of peptides that can be quantified accurately and precisely (Fig. 4F), but that the quantitative suitability for a large number of analytes remained suspicious.

### Extended benchmarking of AvG demonstrates that it produces precise and accurate data

To further evaluate AvG, we created a complex benchmarking set of 4 samples consisting of a mixture of three complex proteomes. The total amount of protein and the proportion of the human proteome was kept constant in all samples, while the proportion of *E. coli* and yeast varied (Table 1, Fig. S9). Six pairwise combinations of the samples are possible, resulting in 12 “ground truth” ratios ranging from 1.2-fold to 10-fold, plus a constant 1:1 ratio of human peptides for all possible comparisons. This experimental design enabled the estimation of reproducibility across many MS runs having different sample compositions, with some compositions more prone to interferences than others.

**Table 1.** Expected ratios of the pairwise relative quantification of samples in the extended benchmarking dataset. From the 4 samples, 6 pairwise combinations can be obtained for relative quantification. This table summarizes the expected ratios for each proteome and each comparison.

| Species | A vs. B | A vs. C | A vs. D | B vs. C | B vs. D | C vs. D |
| --- | --- | --- | --- | --- | --- | --- |
| YEAST | 1 : 4 | 1 : 8 | 1 : 10 | 1 : 2 | 1 : 2.5 | 1 : 1.2 |
| ECOLI | 1.4 : 1 | 3.3 : 1 | 10 : 1 | 2.3 : 1 | 7 : 1 | 3 : 1 |
| HUMAN | 1 : 1 | 1 : 1 | 1 : 1 | 1 : 1 | 1 : 1 | 1 : 1 |

The resulting dataset has a large peptide abundance dynamic range and emulates ratios close to typical thresholds of biological significance for evaluation of DIA analysis tools^11, 20^. We report results at the peptide level (not protein level), as it more accurately reflects the direct measurements made by the mass spectrometer. Protein “roll-up” can mask peptide inaccuracies. We used our DIA benchmarking dataset to evaluate the results of the widely-used mProphet tool compared to data curation with AvG.

The results of the relative quantification accuracy for the baseline mProphet analysis (Fig. S10A) showed high variance and large deviations from the expected values. This phenomenon was readily observed when examining the distribution of ratios for the human peptides, which all should have a nominal ratio of 1:1 in comparing any two samples (Fig. S10A, bottom panel). While these large deviations were more prominent at low intensity, they were observed throughout the entire intensity range. Summarizing across all 6 pairwise comparisons, we observed MAD = 0.20 and *σ*=1.94 around the expected ratio of 1:1. In practical terms, this means that any ratio derived from mProphet results ranging from 0.25:1 to 4:1 (+/ 2*σ*) was not statistically different from 1:1. Relative quantification of the pre-defined ratios for the *E. coli* and yeast peptides was also poor, with a mean percent error of 59% for *E. coli*-derived ratios and 178% for yeast peptides.

After refining the same dataset using AvG, relative quantification accuracy improved dramatically (Fig. S10B). Summary statistics for the human peptides improved to MAD = 0.13 and *σ* =0.27, allowing detection of significant differences at a threshold of 1.54-fold. Relative quantification of the post-defined ratios for the *E. coli* and yeast peptides improved, with a mean percent error of 17% for *E. coli*-derived ratios and 18% for yeast peptides. The FDR calculated after refinement was 0.15% at the measurement level and 0.3% at the peptide level. When scoring mProphet results with AvG metrics, it is clear that many peaks would fall below thresholds that we considered reliable for quantification (Fig. S10C). Improvement in all figures of merit (CV, accuracy, % error, and linear goodness-of-fit) was observed after AvG curation (Fig. S10D). While the total number of peptide analytes dropped after AvG curation, we are confident that these were the most reliable for use in downstream quantitative analyses.

### Performance of AvG under conditions emulating real biological data

Detection of changes in protein levels between two sample classes (e.g., diseased vs. healthy, treated vs. control) is a major paradigm for quantitative proteomics. To evaluate whether AvG curation would help achieve this goal, we simulated biological data to create a realistic scenario in which most peptides in the data set were “unchanged,” while a small minority were up- or down-regulated. This was practically achieved by downsampling the benchmarking data to include 90% human analytes (3000 peptides, unchanged), 5% *E. coli* peptides (positive fold-changes), and 5% yeast peptides (negative fold-changes). The analytes were chosen at random from the larger pool of peptides, allowing us to bootstrap the analysis by selecting different subsets.

We calculated ratios of peptides between the different sample compositions before and after AvG curation, and compared them to the expected ratios (Table 1). Significance (*p*) values were assigned to the ratios using a moderated t-test and corrected for multiple hypothesis testing^21^. Peptides were classified as differentially expressed if their adjusted *p*-value was lower than 0.05 and their absolute fold change was > 2σ of the fold changes for the (unchanged) human peptides present in the downsampled data, and considered “accurate” if their observed ratio was 80-120% of expected. Knowledge of the species-of-origin for each peptide allowed us to classify results as true- or false-positive.

AvG improved the ability to detect changes in protein expression. As an example, we compared sample A to B before and after AvG curation (Fig. 5A). The improved accuracy and precision obtained after AvG resulted in a much higher number of true positive hits (blue and red full disks) and lower number of false positives hits (green full disks not within the grey area). Additionally, the number of accurate measurements (observations between the dashed lines) increased after curation.

**Figure 5:**
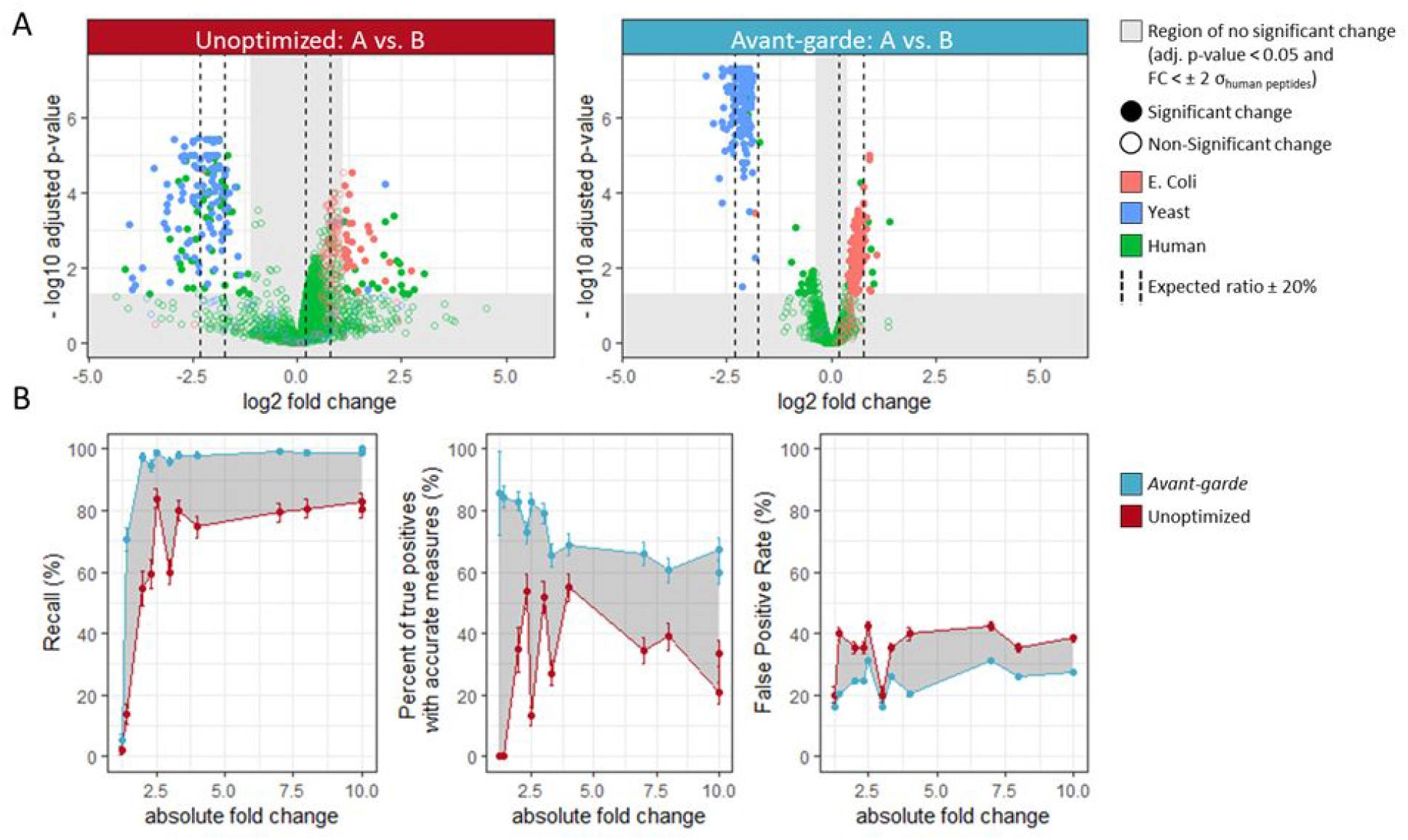
Detection of differentially expressed peptides in unoptimized and curated data. (A) An example of a pairwise comparison (sample A vs. B), with volcano plots of unoptimized (left) and curated (right) data. Each point represents an E. Coli (red), Yeast (blue) or Human (green) peptide. The shaded regions demarcate ranges where detection of differential expression is not statistically viable. The dashed lines represent accuracy boundaries of +/− 20%.(B) Bootstrap (n=1000) analysis of downsampled datasets for recall (sensitivity), accuracy, and false positive rate. Shaded regions indicate improvement in area under the curve after AvG curation. Error bars connote the standard deviation across bootstrap iterations.

To quantify the improvement in performance, we calculated the recall, % of “accurate” measurements, and false positive rates for detection of differentially expressed peptides across the range of fold-changes using 1000 bootstrap iterations as described in Supp. Methods. The results are illustrated in Fig. 5B, with the shaded areas indicating improvement in figures-of-merit achieved after AvG curation. AvG increased the recall and led to a higher number of correct calls of significance for differentially expressed peptides (Fig. 5B, left). After curation, a recall higher than 95% was achieved for any absolute fold change above 2.0. In comparison, unoptimized data could only achieve a recall of ~80% even for 10-fold changes. In addition, we observed an improvement in the percentage of true positive hits that were classified as being *accurate*, with a median improvement of 47% (Fig. 5B, middle). Furthermore, the false positive rate decreased from a median value of 38% to a median of 25%.

The presence of the unchanging proportion of human peptides across all samples allowed us to estimate the minimum detectable fold change (estimated as μ +/− 2σ) before and after data curation. The threshold for uncurated data was 2.3-fold, but fell to 1.4-fold after AvG. Sensitivity (recall) was 70% at this threshold, and 85% of the differential peptides had a calculated ratio within 20% of the true ratio. This demonstrated that we can confidently detect relatively small changes in peptide abundance with properly curated data.

## Discussion

Data curation is an extremely important but often overlooked step in transition-based quantitative proteomics. We have demonstrated that AvG can curate DIA data in an automated manner. AvG tailors the choice of transitions to each dataset to minimize noise and increase the reliability of quantification. A key feature of AvG is that each peptide is reassessed with an independent global scoring module after curation to estimate a dataset-level FDR for *quantitative suitability*, not just detection. Counterintuitively, the number of detected peptides may go down after AvG, but the quality of quantitation for those peptides will be higher. We have empirically demonstrated that the AvG score tends to be low for “decoy” analytes, and data curated with AvG consistently produces FDRs < 1.0% at the thresholds we have defined. Other software tools for DIA analysis typically simply extract the 5-10 most intense transitions from the spectral library. This approach does not guarantee interference-free transitions for analyzing complex biological samples, where curation is paramount.

An alternative approach is to select the transitions that are predicted to be unique to their precursor ion using tools like SRMCollider^22^. However, interferences are difficult to predict due to run-to-run chromatographic variability and changes in sample composition. These tools do not consider retention time or fragment ion intensities, hindering accurate curation. *A priori* prediction of interference, reliant on protein databases and other user choices, does not anticipate real-world LCMS data artifacts. Another *a priori* approach to curation, SWATHProphet^23^, uses spectral libraries embedded with retention time information to anticipate quantitative interferences. In this case, library completeness (again traceable to experimental and user decisions) governs the success of the approach. In contrast, AvG uses an *a posteriori* approach to curate data that explicitly considers LCMS artifacts *and* utilizes prior knowledge but is not limited by it.

SWATHProphet, based on the mProphet discriminant score, also implements an approach for *a posteriori* flagging of poor transitions. Its application requires iterative cycles of optimization and data re-extraction, and again relies on spectral libraries as the primary source of interference detection. Further, it focuses on improving quantitation for peptides that already have a high mProphet score, rather than potentially improving scores of borderline peptides. This approach is apt to produce false negatives and will fail to rescue suitable data signals. AvG focuses on identifying the “cleanest” transitions that are the best suited for quantification, and can improve signal quality for marginal cases. It does not require iterative cycles and is not bound to any specific DIA or PRM workflow, as it is fully implemented as an external tool in Skyline.

The objective of quantitative proteomics is to identify differentially expressed proteins or peptides. Therefore, it is important to evaluate new methods with scenarios that mimic conditions found in real biological milieux. For us, that meant creating a dataset where the majority of peptide analytes were unchanged between two sample classes, while a minority were changing with a known ratio. Moreover, we needed to evaluate ranges of borderline biological significance (1.5 - 2.0-fold) as well as extreme significance (>5-fold) to truly assess method performance. By downsampling and bootstrapping a very large DIA dataset of mixed multi-species proteomes, we could evaluate 1000s of such simulated datasets to test the robustness of AvG. We were pleased to find that AvG could enable discrimination of changes as low as 1.4-fold with fairly high sensitivity and accuracy, and that, across the board, it can add value to the work done by other DIA analysis tools by improving *quantitative suitability* of the data. To us, this means that it can help produce more accurate and reproducible quantification results, providing more granularity in the elucidation of the complex dynamics of proteomes.

AvG’s ensemble-driven scoring strategy is designed to produce very conservative results by penalizing poor-quality signals. Its combined score is a weighted product of run-specific and dataset-wide subscores that intuitively map to common LCMS data quality metrics. AvG penalizes sets of transitions for peptides with any single poorly-scoring metric, making it very sensitive to interferences. However, its evolutionary optimization approach ultimately selects sets of transitions that produce the lowest levels of noise and the highest level of parsimony for signals across the entire dataset. Application of AvG improves selectivity, accuracy, and reproducibility of quantitative DIA proteomics data. The resulting curated data is comparable to the current gold-standard of expert human curation, but obtainable in a fraction of the time. AvG’s compatibility with a variety of acquisition modes (DIA or PRM), data sources (e.g. Orbitrap and TOF), upstream DIA identification tools (e.g. EncyclopeDIA, Specter, mProphet, etc.), and Skyline integration should make it attractive for broad utilization in the field.

## Acknowledgements

The authors would like to thank Nicholas Pythoud, Joanna Bons, Alexandre Burel and Christine Carapito for beta-testing the software. This work was funded by U54 HG008097 to JDJ.

## Author Contributions

A.S.V.J. conceived of the idea, designed and performed experiments, collected the data, authored software, and wrote the manuscript. R.P. provided help with the data analysis of the benchmarking datasets. N.S. and B.M. adapted Skyline to facilitate the use of Avant-garde as an External tool. K.K. provided help with the statistical analysis of the benchmarking datasets. K.C.D. and A.O. carried out experiments and collected data for the P100 dataset, and beta-tested the software. M.J.M. and S.A.C. provided laboratory resources and guidance on the manuscript. J.D.J. provided laboratory resources, provided input on the software, provided guidance on the experimental design and wrote the manuscript.

## Code availability

Avant-garde DIA is an open-source software tool available as an R package and as an Skyline External tool at https://github.com/SebVaca/Avant_garde. Avant-garde can be directly downloaded from tool Store interface within Skyline or from the Skyline tool Store at https://skyline.ms/skyts/home/software/Skyline/tools/details.view?name=AvantGardeDIA.

## Data availability

The data that support the findings of this study are available from the corresponding author upon request.

