## Supplemental Materials for "*Avant-garde*: An automated data-driven DIA data curation tool"

### Supplementary figures

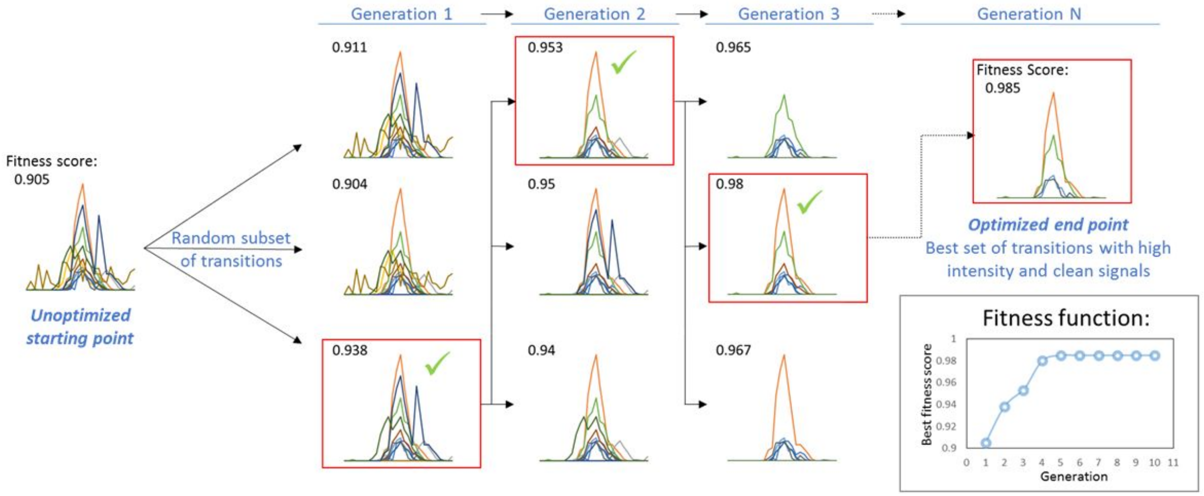


***Supp. Figure 1:*** Principles of "natural selection" of transitions. The transition refinement tool uses a genetic algorithm. For each generation, a population of subsets of transitions is randomly selected and scored with a fitness function. Here we illustrated a population of 3 solutions. The best-scoring solution is kept and is used as the starting point for the next generation. Over successive steps, the population "evolves" towards an optimal solution, as shown in the inset panel showing the highest score of the fitness function in each generation. The set of transitions obtaining the highest score will be most suitable for quantification.

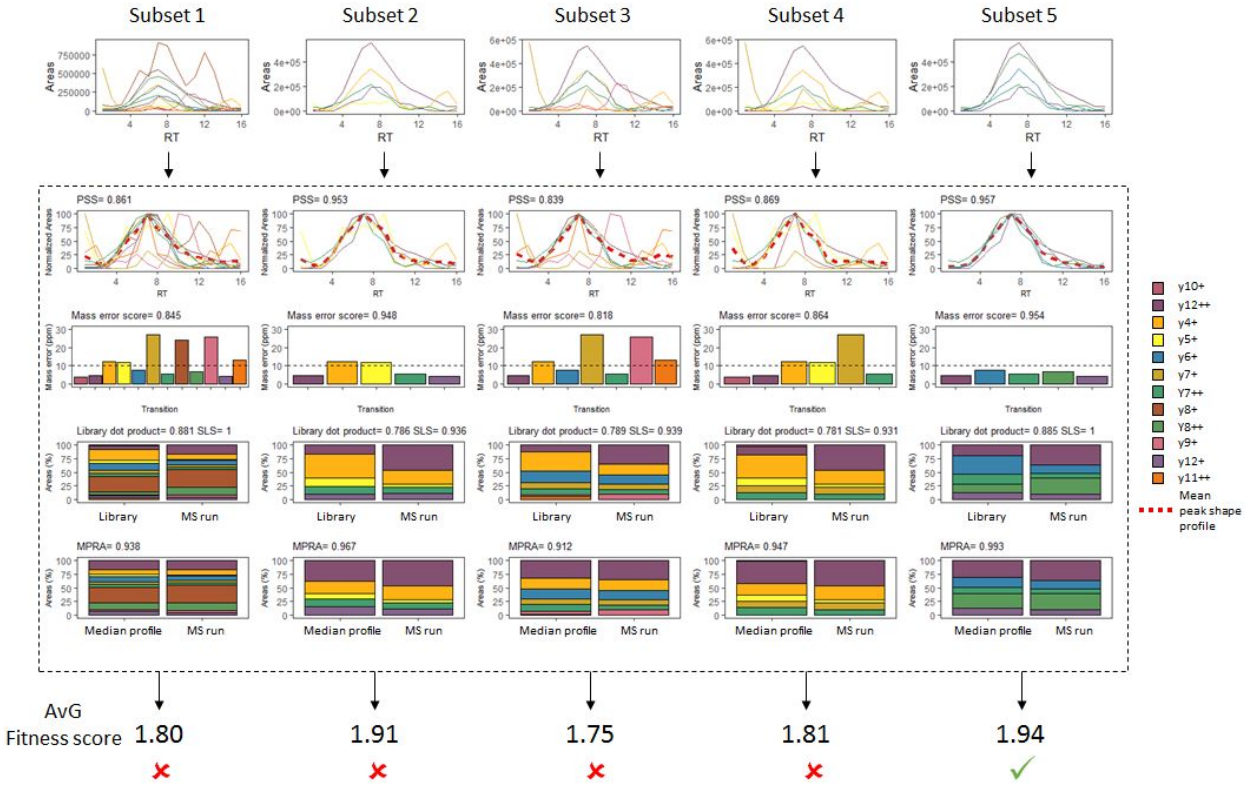


***Supp. Figure 2:*** <u>Avant-garde uses a fitness score that combines run-specific and dataset-wide features.</u> The fitness function principle is illustrated here for peptide TPSIQPSLLPHAAPFAK. Twelve transitions were extracted for this peptide. In this example five subsets of transitions for this peptide are scored and compared in order to determine which one is the most suited for its quantification. The fitness function of the genetic algorithm is a combination of run-specific and dataset-wide subscores. The peak shape similarity score (PSS) evaluates the similarity of each transition to all others transitions used to quantify a peptide by comparison to a mean peak shape profile (dashed red line). The mass error score penalizes sets of transitions with any large deviations from their expected masses. The spectral library similarity score (SLS) evaluates the similarity between DIA peak areas and spectral library intensities. The median profile of relative areas (MPRA) score is a dataset-wide score that measures how similar the signals in a given run are to all other runs. This metric provides an ensemble-driven element that captures the similarity of the signals across the entire dataset in the composite *AvG fitness score*. Subset 5 would be chosen here as all the transitions in the set have similar shapes, low mass error, and the relative areas match the best with the spectral library and with all other runs in the entire dataset.

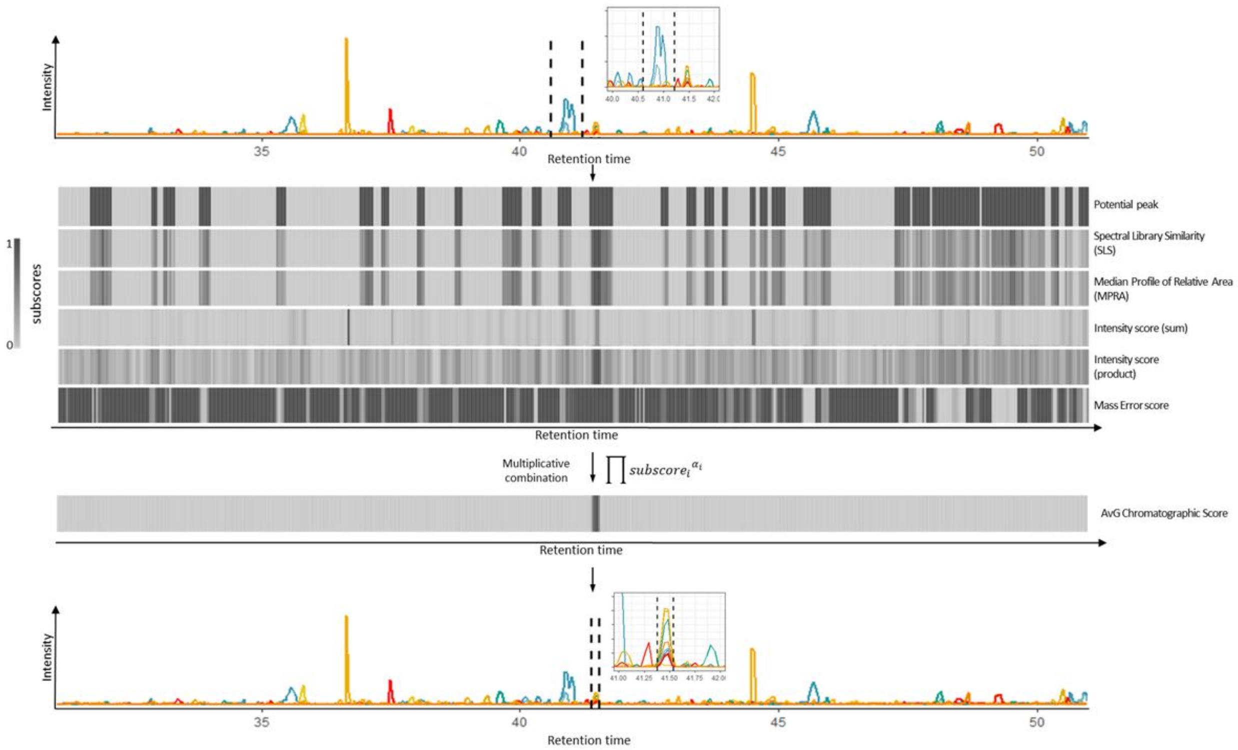


<u>Supp. Figure 3: The peak boundaries refinement tool is effective in wide search windows.</u> Detection of a chromatographic peak using the *AvG chromatogram score* in a 20-minute search window. Data shown are expanded from the example shown in Fig. 1. AvG refines peak integration boundaries by calculating chromatographic subscores at each time point in the raw data, and combines them as a weighted product (*AvG chromatographic score*). The maximum value of this score corresponds to the most likely retention time of the analyte. On the top panel an example of a wrong peak picking is shown. After AvG peak boundary correction the correct peak is integrated (bottom panel). AvG does not require spiked-in retention time peptides as this module performs well even when working on wide chromatogram search windows.

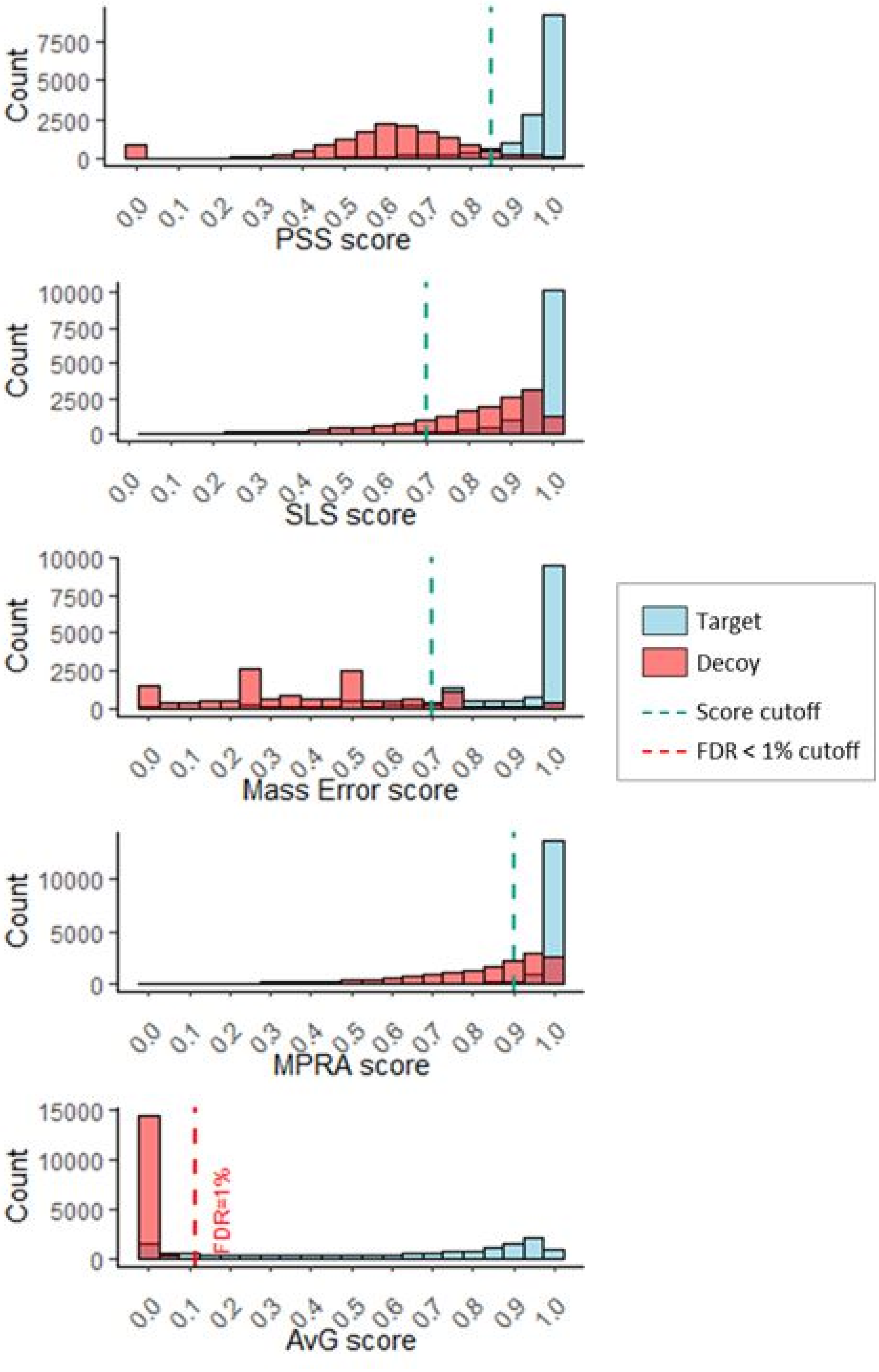


Supp. Figure 4: Target-decoy distribution of subscores and the AvG score shows good discrimination between low- and high-quality signals. Histograms showing the target-decoy distribution for subscores and the AvG score. The green and red vertical lines correspond to the individual score threshold for each subscore and the AvG score threshold to guarantee an FDR below 1%. The data are shown at the measurement level, corresponding to one peptide in one run.

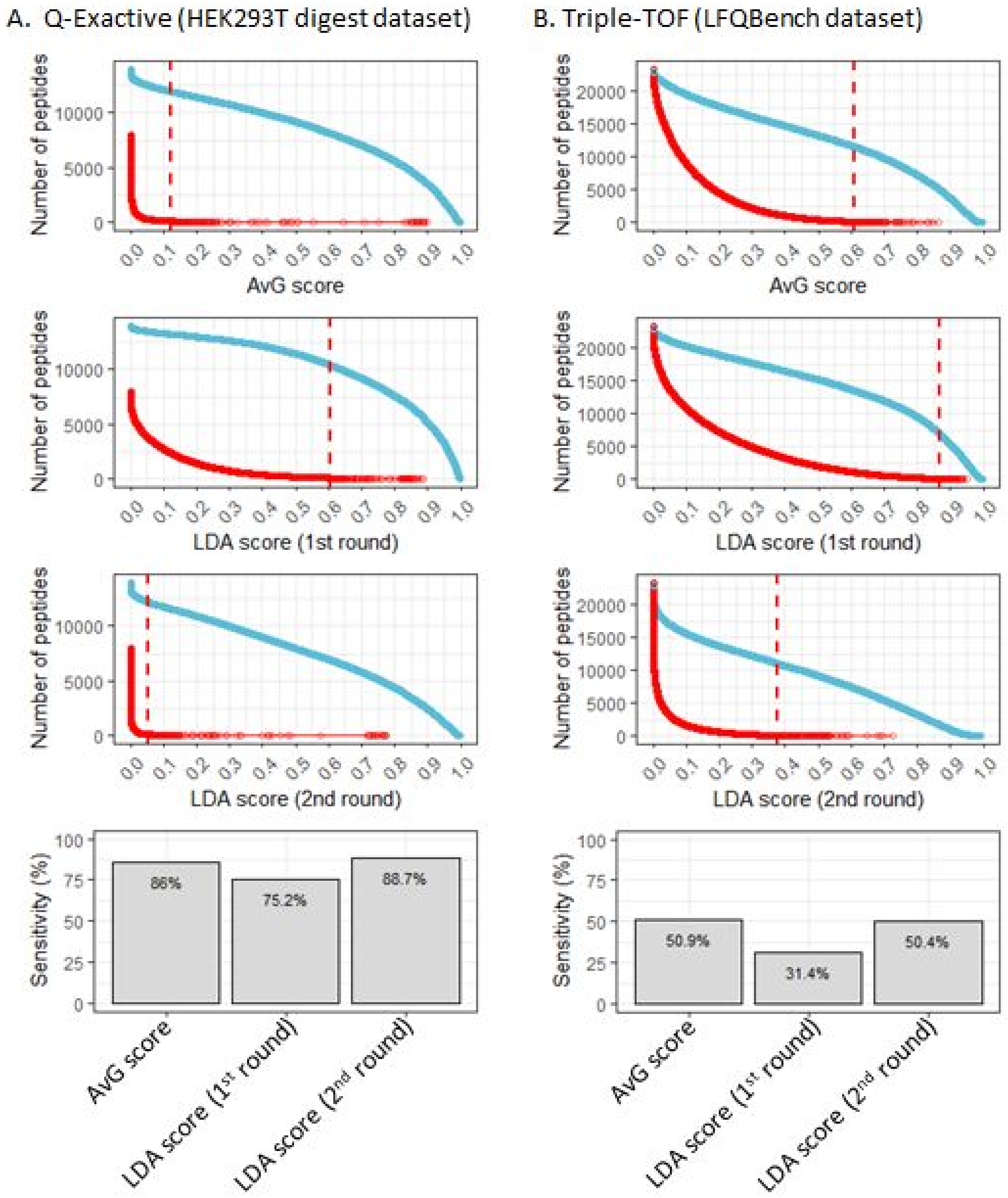


Supp. Figure 5: Comparison of empirical vs. linear discriminant analysis-determined (LDA) AvG weighting. Examples demonstrate the score cutoff determined by each method to obtain a 1% FDR for quantitative suitability. AvG score uses empirically determined weights, while the LDA scores use LDA-determined weights. Two rounds of LDA optimization were required for technical reasons (see Supp. Methods). Blue: cumulative number of target peptides. Red: cumulative number of decoy peptides. Final sensitivity values (bottom row) are comparable for both Orbitrap (A) and Triple-TOF (B) data.

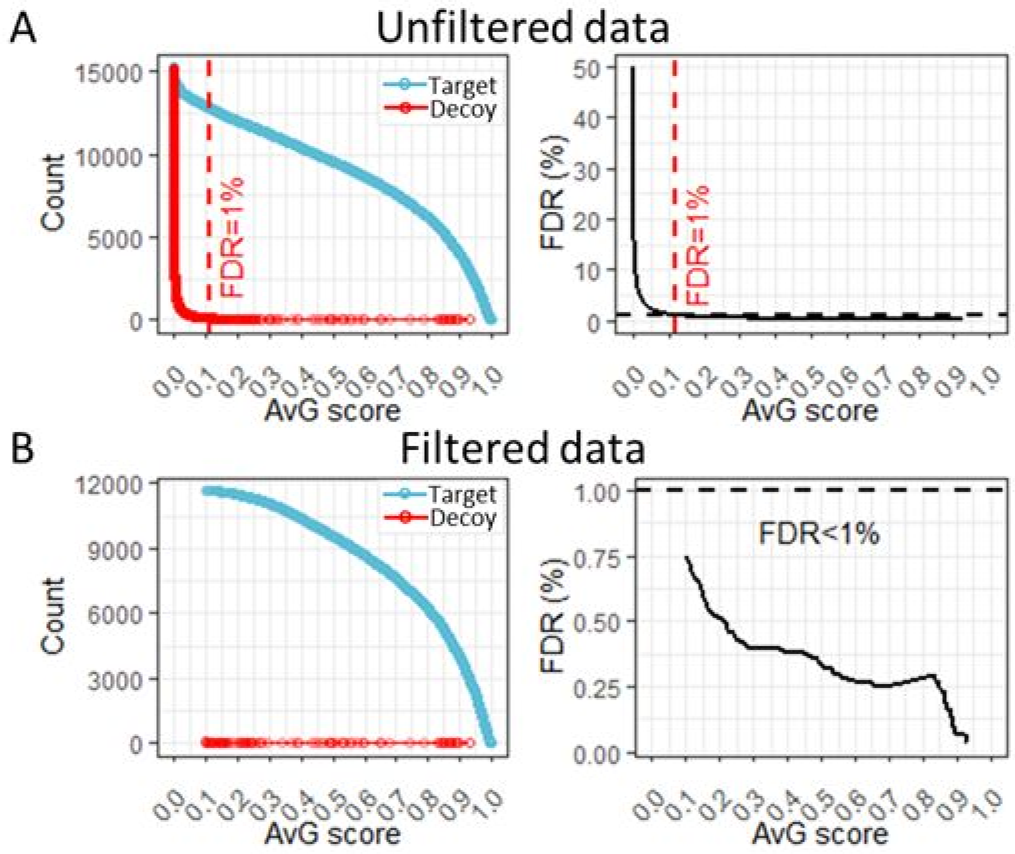


<u>Supp. Figure 6: FDR evaluation in HEK293T whole cell digest sample after signal refinement by AvG</u>. Counts of target and decoy measurements and FDR as a function of the AvG score. The data was unfiltered (A) or filtered (B) before evaluating the FDR. The filtering thresholds applied to the subscores, to guarantee a minimal level of signal quality, were: SLS >0.7, mass error score>0.7, PSS >0.85, MPRA > 0.9, and AvG score >0.1.

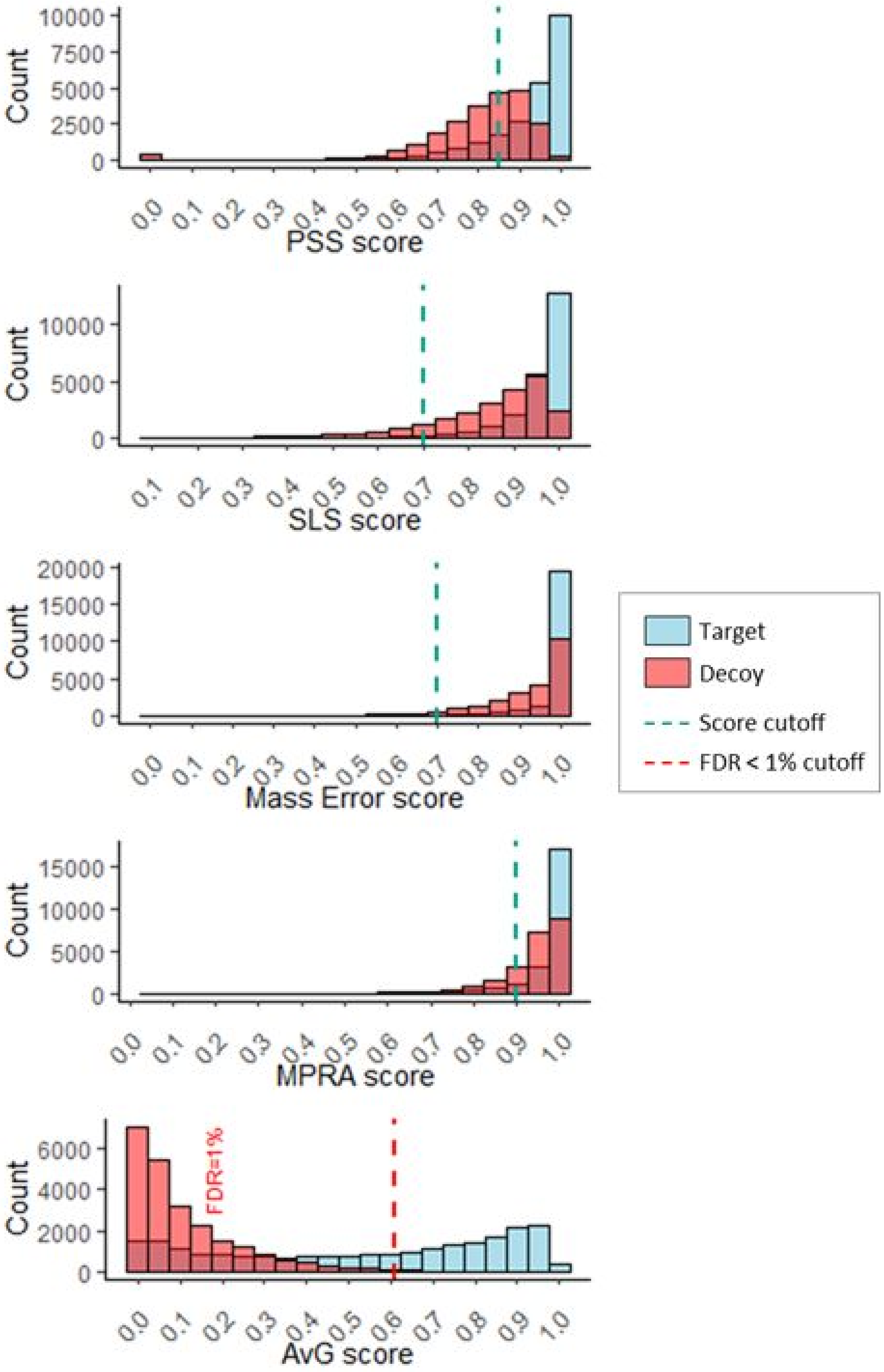


Supp. Figure 7: Target-decoy distribution of subscores and the AvG score for Q-TOF data. Histograms of the target and decoy distributions. The green and red vertical lines correspond to the individual score threshold for each subscore and the AvG score threshold to guarantee an FDR below 1% (AvG score<0.61). The data is shown at the measurement level, corresponding to one peptide in one run.

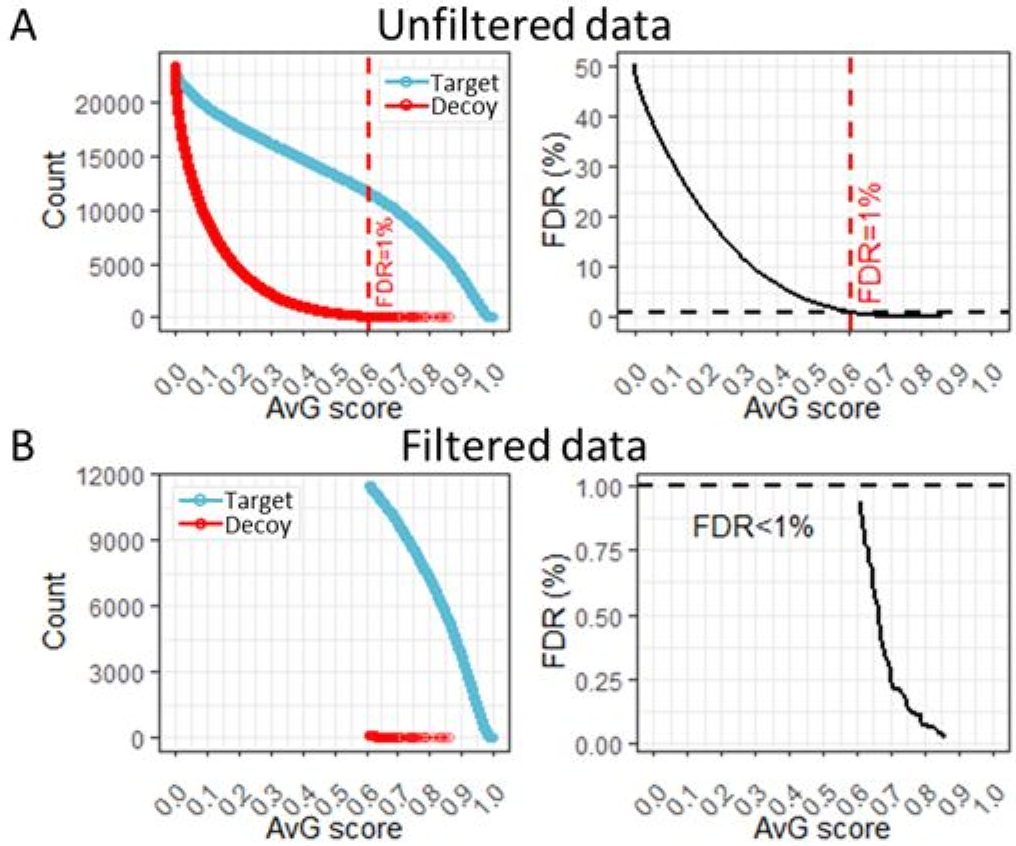


Supp. Figure 8: False discovery rate evaluation for the LFQBench dataset after signal refinement by AvG. Counts of target and decoy measurements and FDR as a function of the AvG score. The data was unfiltered (A) or filtered (B) before evaluating the FDR. The filtering thresholds applied to the subscores, to guarantee a minimal level of signal quality, were: SLS >0.7, mass error score>0.7, PSS >0.85, MPRA > 0.9, and AvG score >0.61.

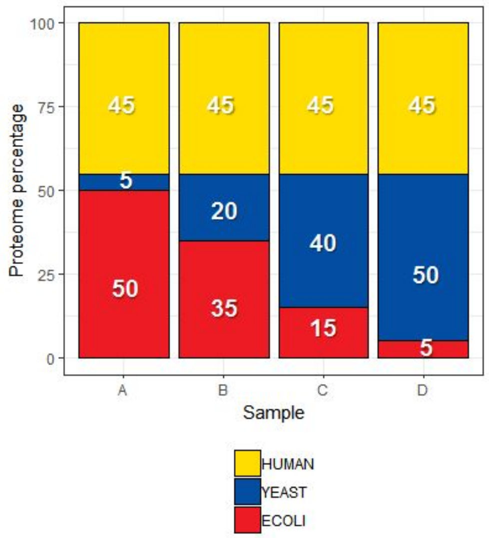


Supp. Figure 9: Extended benchmarking DIA dataset design. The benchmarking dataset consists of 4 hybrid proteome samples. Each one is a mixture of three complex proteomes in different proportions. In all four samples, the total amount of protein is kept constant. The proportion of the human proteome is the same for all samples. The proportion of the yeast and *E. coli* proteomes changes from one sample to another. This dataset was designed to take advantage of the 6 sample pairwise combinations that generate 12 expected peptide ratios for *E. coli* and yeast showed in Table 1.

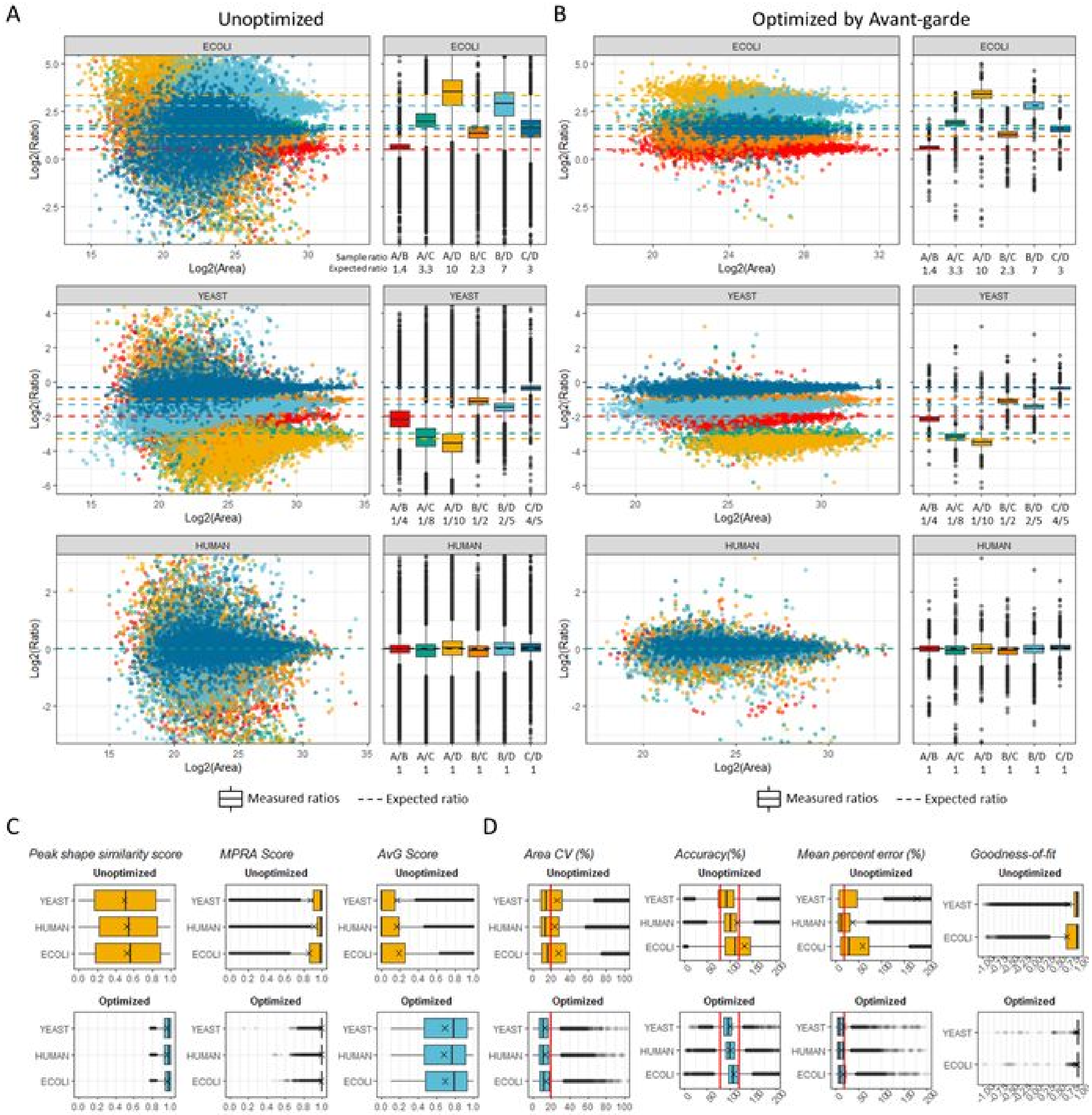


Supp. Figure 10: Relative quantification and distribution of ratios for extended benchmarking. Unoptimized dataset (A, Skyline/mProphet data filtered with q-value<0.01) and AvG-curated dataset (B). Each dot represents a ratio calculated for a given peptide in a given run. The dashed lines represent the expected ratios. The boxplots show the distribution of the observed ratio of each pairwise combination. The box plot elements are: center line, median; box limits, upper and lower quartiles; whiskers, 1.5x interquartile range; points, outliers. (C) Distribution of selected AvG subscores calculated pre- (top) and post-curation (bottom). (D) Metrics to evaluate the accuracy and precision (CV) of the relative quantification pre- (top) and post-curation (bottom). The red vertical lines demarcate thresholds commonly used in quantitative proteomic studies: CV and % error < 20%, accuracy boundaries of +/- 20%. Goodness-of-fit was evaluated by the correlation between the 6 observed and expected ratios for each proteome. Crosses show the mean value of each metric.

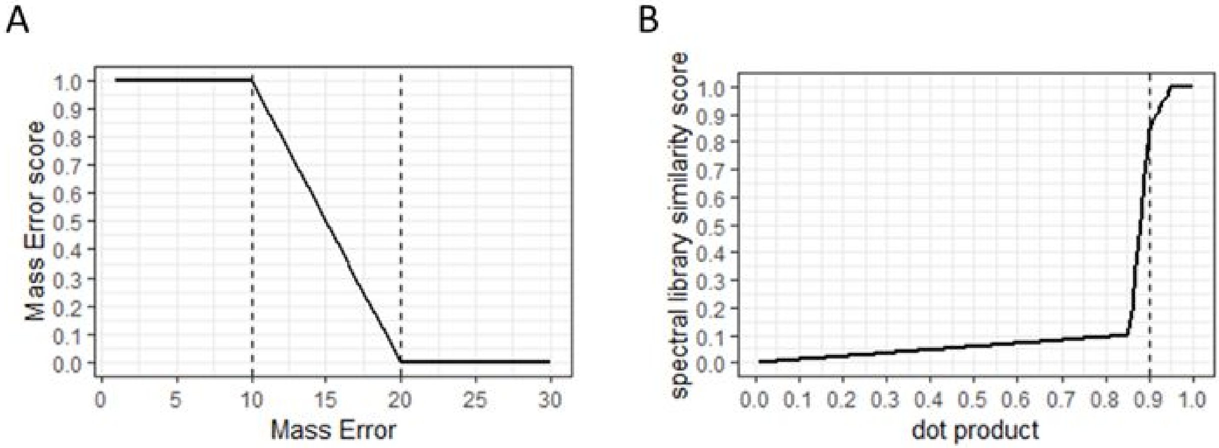


Supp. Figure 11: Transformations used during AvG scoring (A) Transformation of the mass error in ppm into the Mass Error score used by AvG. This example shows the case where the user-defined parameters for the mass error are a mass error tolerance equal to 10 ppm and a mass error cut off equal to 20 ppm. (B) Transformation of the dot product between the measured areas under the curves and the corresponding intensities in the spectral library, into the spectral library similarity score (SLS score). This example shows the case where the user-defined parameter for the dot-product limit is equal to 0.9.

**Supplementary methods**

<u>HEK293T cell digest</u>

HEK293T cells were cultured in DMEM (Gibco; 11995) supplemented with 10% heat-inactivated FBS (Sigma; F4135). Once cells reached ~95% confluence they were harvested by scraping. Cells were pelleted at 1,000g for 2 min. The supernatant was then removed, and the cell pellet was frozen in liquid nitrogen. HEK293T cells were lysed by 5 min of exposure on ice to a lysis buffer (8 M urea, 75 mM NaCl, 50 mM Tris-HCl, pH 8.0, 1 mM EDTA, 2 µg/mL aprotinin (Sigma; A6103), 10 µg/mL leupeptin (Roche; 11017101001), 1 mM PMSF (Sigma; 78830)). The sample was centrifuged for 10 min at 20,000g. The protein concentration of HEK293T proteins was determined by BCA assay to be 4.3 µg/µl. 10 mg of protein was reduced (5 mM dithiothreitol, 45 min) and alkylated (10 mM iodoacetamide, 45 min). A Tris-HCl solution (50 mM, pH 8) was used to dilute the samples by a factor of 4 to reach a concentration of 2 M urea. A two-step digestion protocol was used to digest the lysate: Lys-C was used in a 1:50 enzyme-to-substrate ratio (Wako Chemicals; 129-02541) for 2 h at 30 °C, then the lysate was digested overnight at room temperature with trypsin in a 1:50 enzyme-to-substrate ratio (Promega; V511X) on a shaker. Formic acid (FA; 0.5% final concentration) was added to stop the digestion. The sample was split into four aliquots and loaded onto four 100-mg-capacity C18 Sep-Pak cartridges (Waters) for desalting. The four aliquots were eluted with 50% acetonitrile (ACN)/0.1% FA, pooled together, and vacuum-concentrated to dryness.

<u>E.Coli Digest</u>

DH5α E. coli were grown in Luria broth at 37 °C overnight. Cells were pelleted by centrifugation, washed once with cold PBS, flash-frozen in liquid nitrogen, and stored at –80 °C until processing. For generation of the E. coli lysate digest, the cell pellet was thawed on ice. Lysozyme (Sigma) was added to the thawed pellet, and the mixture was placed on ice with periodic vortexing until viscous. The cells were resuspended in 8 M urea, 50 mM ammonium bicarbonate plus protease inhibitors (Roche), and the solution was sonicated with a probe sonicator for 2 min, 3 s on, 2 s off, until no longer viscous. After centrifugation at 15,000g for 30 min at 4 °C, protein concentration was measured by Bradford assay (Bio-Rad). Disulfide bridges were reduced (10 mM TCEP (tris(2-carboxyethyl)phosphine), Thermo) and alkylated (10 mM iodoacetamide, Thermo; 30 min; room temperature; in the dark). The lysate was diluted to 1.5 M urea with ammonium bicarbonate (50 mM) and digested overnight with a trypsin-to-substrate ratio of 1:100. The digest was desalted on C18 Sep-Pak cartridges (Waters). After vacuum centrifugation, dried peptides were resuspended to 1 mg/mL in 30% ACN/0.1% FA and stored at –80 °C.

<u>Extended benchmarking DIA dataset</u>

Mass spec-compatible Yeast digest was purchased from Promega. The E. Coli, yeast and HEK293T digest were resuspended to 1 mg/mL in 30% ACN/0.1% FA. Four samples, with sample composition described in Supp. Fig. 9, were generated for the benchmarking dataset. A fifth sample that had the same quantity of protein of each proteome was also generated to obtain the spectral library. To generate the samples, the volume of each proteome digest corresponding to the desired protein quantity were mixed together, vacuum-concentrated to dryness, and resuspended to 500 ng/ul in 5% ACN/0.1%TFA. The 4 samples for the DIA benchmarking dataset were analyzed in DIA mode in 4 replicates.

Calibration curve in HEK293T cell digest

The HEK293T digest was resuspended using 0.1% TFA, and a mixture containing 95 synthetic peptides at known individual concentrations was spiked into it to generate a five-point calibration curve. Each point was designed to contain 1 µg of HEK293T digest and 6.75, 13.5, 27, 54, or 108 ng of total amount of peptide on the column. The 5 samples were analyzed in DIA mode in triplicates.

We generated a spectral library by first searching the DDA runs with Spectrum Mill v. B.06.01.201 using a FASTA containing the 95 synthetic peptide sequences and the UniProt human protein sequences (version dated 17 October 2014). Results were auto-validated to a false discovery rate of 1%, exported as a PepXML search result file, and loaded into Skyline to generate a spectral library in blib format.

The data analysis was performed using Skyline. Eleven unmodified peptides from the HEK293T background were chosen as standards to calibrate the retention times. The transition selection and peak boundaries was performed by AvG. The coefficient of variation were calculated on the triplicate measurement of the area of each analyte. The percent error was obtained by calculating the concentrations of each calibration point with the linear regression equation found for each peptide. The percent error was calculated as the absolute value of the difference between the measured and the expected concentration over the expected concentration.

P100 dataset:

The P100 samples were prepared exactly as described in Abelin et al. 2016[1]. In short, cells were cultivated, perturbed with 32 drugs each with 3 biological replicates. Cells were lysed for 30 min at 4 °C in lysis urea buffer (8 m urea; 75 mm NaCl, 50 mm Tris HCl pH 8.0, 1 mm EDTA, 2 µg/ml aprotinin (Sigma), 10 µg/ml leupeptin (Roche), 1 mm PMSF (Sigma), 10 mm NaF, Phosphatase Inhibitor Mixture 2 (1:100, Sigma), Phosphatase Inhibitor Mixture 3 (1:100, Sigma). Lysates were centrifuged at 15,000 × g for 15 min. Protein concentrations were measured (660 protein assay, Pierce). Reduction, Alkylation and digestion were performed on a Bravo robotic liquid handling platform (Agilent). Five hundred micrograms of protein were used for reduction in 100 mm DTT, alkylation in 200 mm IAA, dilution to 2 m urea in 50 mm Tris (pH 8.0), and digestion with 0.5 µg/µl (1:50) sequencing-grade modified trypsin (Promega) in 400µl volumes per sample at 37 °C overnight. Digestion was stopped by bringing the samples to a final concentration of 0.5% TFA. Acidified samples were loaded onto a 25 mg capacity C18 SepPak in a 96-well plate format (Waters) for desalting. Samples were eluted using 50% ACN/0.1% trifluoroacetic acid and vacuum concentrated to dryness. Then the samples were phospho-enriched pn an AssayMAP Bravo robotic system (Agilent). The desalted samples were reconstituted in 80% ACN/0.1% TFA. Prior to sample loading the Agilent AssayMAP Fe-(III)-NTA cartridges were washed with water, stripped with 100 mm EDTA, and loaded with 100 mm FeCl3. Fe-(III)-NTA cartridges were primed with 1:1:1 ACN/methanol/0.01% acetic acid. Samples were loaded at 20 µl/min and flow-throughs were re-loaded onto cartridges eight additional times. Cartridges were washed with 80% ACN/0.1% TFA, and peptides were eluted with 500 mm K2HPO4 (pH7) at 5 µl/min. Eluates were vacuum concentrated to dryness, and subsequently desalted using AssayMAP RP-S cartridges according to the manufacturer's instructions. The 96 samples were analyzed in DIA mode.

For the data analysis we focused on the 95 phosphopeptides that constitute the P100 assay[1], which had isotopically labelled heavy peptide counterparts spiked into the sample. The dataset was analyzed using Skyline. For the manual validation: 3 to 5 transitions per peptide were extracted. The transitions were the ones chosen for the P100 assay. The data were visually inspected, interfered transitions were removed, and peak boundaries manually corrected by an expert in the field. For the unoptimized dataset: the 5 most intense transitions from the spectral library were chosen and the peak boundaries were defined by Skyline. For the optimized version, all possible transitions (b,y above b4 and y4) were extracted and the transition selection and peak boundaries was performed by AvG.

Evaluation of AvG using the LFQBench dataset

The LFQBench data[2] was downloaded from ProteomeXchange (dataset PXD002952). We focused on the HYE110 dataset analyzed with 64 variable isolation width windows on an AB Sciex Triple-TOF 6600. The raw WIFF files were imported into Skyline Daily (v.4.1.1.18118). We used the spectral library provided by the study's authors (ecolihumanyeast_concat_mayu_IRR_cons_openswath_64w_var_curated.csv) that consisted of precursors with only 6 annotated fragment ions in CSV format compatible with OpenSWATH.

For the optimized dataset the retention time standard peptides (iRTs, Byognosis) were used to predict the expected elution times. Skyline signal extraction parameters were: 30000 resolving power for MS/MS, chromatograms were extracted 5 minutes around the predicted RT, all 6 six ions were extracted for each precursor.

The results from Avant-garde (peak boundaries and the selected transitions) were imported back into the same Skyline file.

Evaluation of AvG using our extended benchmarking DIA dataset

To build the spectral library a sample constitute of an equal protein amount of the three proteomes was analyzed. This sample was analyzed using gas phase fractionation using 12 MS runs[3]. Each injection was analyzed with narrow-window DIA where the instrument cycles through 25 2-m/z DIA windows and focuses only on 50 m/z at the MS1 level per MS run. Twelve injections are necessary to systematically and comprehensively monitor the 400 to 900 m/z range with narrow windows. The data were searched with SpectrumMill against a merged database of the human, E. Coli and yeast database. The search was done with large tolerance at the MS1 level (1 m/z) and small tolerance at the MS2 level 10ppm. The search results were validated using Percolator 3.0 at a false discovery rate lower than 1%. The list of validated spectra was imported into Spectrum Mill, a PepXML search result file was generated and loaded into Skyline to generate a spectral library in blib format.

Eighteen thousand peptides (6,000 of each proteome) were randomly chosen from the spectral library and extracted using Skyline. A subset of 15 unmodified human peptides whose retention times were distributed across the chromatographic gradient were chosen as retention time standard peptides.

For the unoptimized dataset the chromatograms were extracted using Skyline and validated using Skyline's implementation of mProphet. The 6 most intense transitions from the spectral library were extracted and a mProphet model was trained using the corresponding 18 thousand decoy peptides. Only the peaks with a q-value lower than 0.01 were reintegrated.

For the optimized dataset the chromatograms were extracted using Skyline. The 10 most intense transitions for each peptide were extracted and the peak boundaries were determined by Skyline, which was using our 15 RT standard peptides for retention time prediction. The data was refined a posteriori by AvG to select at least 4 transitions per peptide and correct the peak boundaries. The data was scored and filtered to less than 1% FDR (spectral library similarity >0.7, mass error score>0.7, peak shape similarity score >0.85, MPRA score> 0.9 and AvG Score>0.1).

In order to simulate "real life" biological samples, we downsampled the dataset by randomly selecting a 3,000 human peptides, and then randomly selecting a lower number of peptides for yeast and E.Coli that represented 5% percent of the total number of human peptides. For each downsampled dataset, a two-tailed two-sample moderated t-test from the limma R-package were calculated for the quadruplicate log2 transformed areas[4]. The p-values were adjusted for multiple hypothesis testing using the Benjamini-Hochberg method. The peptides were classified as significantly differentially expressed (positive hits) if their adjusted p-value was lower than 0.05 and their absolute fold change was higher than 2 times the standard deviation of the fold changes for the human peptides. Peptides were classified as not differentially expressed (negative hits) if either the adjusted p-value was lower than 0.05 or the absolute fold change was lower than 2 times the standard deviation of the fold change for the human peptides. Peptides were classified as accurate if their observed ratio was in the 80-120% range of the expected value. The recall for the detection for differentially expressed peptides was estimated here by the number of true positive hits over the total number of differentially expressed peptides, i.e. yeast and E.Coli peptides (Recall= TP/(Total number of yeast and E.Coli peptides in the downsampled dataset)). The false positive rate (FPR) was determined by calculating the number of false positive hits over the total number of proteins found to be differentially expressed (FP/(TP + FP)). The process described above (downsampling, statistical testing and performance evaluation) was iterated a thousand times to reduce the effect of outlier in the evaluation of the performances of the unoptimized and the optimized dataset by AvG.

LC-MS method

Overlap DIA method on Q-Exactive HF+ (used for P100 dataset and calibration curve spiked in HEK 293T digest):

The P100 dataset and the calibration curve spiked in HEK 293T digest were analyzed with an Orbitrap Q-Exactive HF Plus (Thermo Fisher Scientific) mass spectrometer coupled to a nanoflow Proxeon EASY-nLC 1000 UHPLC system (Thermo Fisher Scientific). The mass spectrometer was used in positive mode and was equipped with a nanoflow ionization source (James A. Hill Instrument Services, Arlington, MA); the spray voltage was set at 2.00 kV. The LC system, the column, and the electrospray voltage source (platinum wire) were connected via a stainless steel cross (360 µm; IDEX Health & Science; UH-906x). The column was heated to 50 °C. A volume of 3 µl was injected onto an in-house packed 20 cm × 75 µm diameter C18 silica picofrit capillary column (1.9-µm ReproSil-Pur C18-AQ beads, Dr. Maisch GmbH, r119.aq; Picofrit 10-µm tip opening, New Objective, PF360-75-10-N-5). The mobile phase had a flow rate of 250 nL/min and consisted of 3% ACN/0.1% FA (solvent A) and 90% ACN/0.1% FA (solvent B). The column was conditioned before each sample injection. Peptides were separated using the following LC gradient: 0–3% B in 3 min, 5–40% B in 50 min, 40–90% B in 1 min, stay at 90% B for 5.5 min, and 90–50% B in 30 s. DDA and DIA data were acquired on the same instrument. For the MS1 scans, the resolution was set at 60,000 at 200 m/z and the automatic gain control (AGC) target was $3 \times 10^6$ with a maximum inject fill time of 20 ms. For DDA, MS2 scans on the top 12 peaks doubly charged and above were acquired at a resolution of 15,000, AGC target of $5 \times 10^4$ with maximum inject fill time of 50 ms. Isolation widths were set to 1.5 m/z with a 0.3 m/z offset. The normalized collision energy (NCE) was set to 27 and dynamic exclusion was set to 10 s. For DIA, an overlap DIA method was used with 56 × 22 m/z isolation windows covering the 400–1,000 m/z range. In this method, the isolation windows in two consecutive cycles have an offset of 11 m/z. The default charge state was 4, the resolution was 30,000 at 200 m/z, the AGC target was $1 \times 10^6$, the maximum inject fill time was 50 ms, the loop count was 27 and the NCE was set to 27.

Overlap DIA and narrow-window DIA method on Q-Exactive HFX (used for the extended benchmarking DIA dataset):

The extended benchmarking DIA dataset was analyzed with an Orbitrap Q-Exactive HFX (Thermo Fisher Scientific) mass spectrometer coupled to a nanoflow Proxeon EASY-nLC 1200 UHPLC system (Thermo Fisher Scientific). The mass spectrometer was used in positive mode and was equipped with a nanoflow ionization source (James A. Hill Instrument Services, Arlington, MA); the spray voltage was set at 2.00 kV. The LC system, the column, and the electrospray voltage source (platinum wire) were connected via a stainless steel cross (360 µm; IDEX Health & Science; UH-906x). The column was heated to 50 °C. A volume equivalent to 500ng of protein on column was injected onto an in-house packed 20 cm × 75 µm diameter C18 silica picofrit capillary column (1.9-µm ReproSil-Pur C18-AQ beads, Dr. Maisch GmbH, r119.aq; Picofrit 10-µm tip opening, New Objective, PF360-75-10-N-5). The mobile phase had a flow rate

of 200 nL/min and consisted of 3% ACN/0.1% FA (solvent A) and 90% ACN/0.1% FA (solvent B). The column was conditioned before each sample injection. Peptides were separated using the following LC gradient: 2–6% B in 1 min, 6–30% B in 74.5 min, 30–60% B in 7.5min, 60–90% B in 1 min, stay at 90% B for 5 min, and 90–50% B in 2min.
For the MS1 scans, the resolution was set at 60,000 at 200 m/z and the automatic gain control (AGC) target was $3 \times 10^6$ with a maximum inject fill time of 20 ms.
For DIA, an overlap DIA method was used with 68× 18 m/z isolation windows covering the 400–1,000 m/z range. In this method, the isolation windows in two consecutive cycles have an offset of 9 m/z. The default charge state was 4, the resolution was 15,000 at 200 m/z, the AGC target was $1 \times 10^6$, the maximum inject fill time was 18ms, the loop count was 34 and the collision energy was fixed for each window using the following equation: CE=0.0459 m/z - $6x10^{-5}$. The m/z value corresponded to the center of each DIA window.
For narrow-window DIA, we used 12 different instrument methods. Each one used 25× 2 m/z non-overlapped isolation windows covering 50 m/z range. Together the 12 MS runs covered the 400-1000 m/z range. The default charge state was 4, the resolution was 15,000 at 200 m/z, the AGC target was $1 \times 10^6$, the maximum inject fill time was 25ms, the loop count was 25 and the collision energy was fixed for each window using the equation described on the paragraph above.

<u>Calculation of AvG subscores</u>

- Peak shape similarity (PSS) score:

For a given peptide and a given subset of transitions, the similarity score shows the resemblance of the peak shape of a given transition to all the others transitions in the subset. This score is calculated by first normalizing each transition to the maximum intensity value within the integration boundaries. By doing this, the peak shape comparison is not dependent on the intensity of each transition but only on its shape. A mean peak shape profile is then created by calculating the mean of the normalized intensities for all transitions at each time point. The mean was chosen in order to better reflect the presence of interferences would be reflected and avoid smoothing them away. This will amplify the difference of peak shapes when a interference is present.
The similarity is determined by calculating the mean of all the dot products, calculated for each transition, of the normalized intensities of each transition and the mean profile.
To make sure that the similarity score is not influenced by just a single highly-scored transition, a second mean is calculated after removing the transition with the highest dot product value. This penalizes even further the set of transitions where an interference is present, ensuring that only the set of transitions that have similar peak shapes obtain a high score. The mean of these two dot product values corresponds to the peak shape similarity score.

For a given peptide using a given set of transitions in a given MS run:

$n = total\ number\ of\ transitions\ in\ the\ set$

$u_i = vector\ of\ the\ intensities\ of\ a\ transition\ i\ in\ elution\ time\ order$

$v$ = vector of the intensities of the mean peak shape profile in elution time order
$k$ = Index of the transition for which the normalized dot product is the highest
$K$ = set of all indices from 1 to $n$ except $k$

$$\text{Peak Shape Similarity score} = \frac{\sum_{i=1}^{n} \frac{u_i \cdot v}{\|u_i \cdot v\|}}{n} \times \frac{\sum_{i \varepsilon K} \frac{u_i \cdot v}{\|u_i \cdot v\|}}{n-1}$$

- Mass error score:

The mass error score is a mean of the mass error measured at each chromatographic point between the integration boundaries weighted by the intensity. The user defines a tolerance threshold in ppm below which the score is equal to 1 and a cut-off threshold above which the score is equal to 0 (Fig. S11A). The mass error score is defined as follows:

For a given peptide using a given set of transitions in a given MS run:
$m_{measured}$ = absolute value of the mass error measured at a given chromatographic point (in ppm)
$m_{tol}$ = mass error tolerance (in ppm)
$m_{cutoff}$ = mass error cutoff (in ppm)

$$\text{mass error score} = 1 \text{ if } m_{measured} \leq m_{tol}$$
$$\text{mass error score} = \frac{m_{cutoff}}{(m_{cutoff} - m_{tol})} + \frac{m_{measured}}{(m_{tol} - m_{cutoff})} \text{ if } m_{tol} < m_{measured} \leq m_{cutoff}$$
$$\text{mass error score} = 0 \text{ if } m_{measured} > m_{cutoff}$$

- Mean profile of relative areas (MPRA) score

For a given peptide and a given subset of transitions, the area under the curve of each transition is normalized to the sum of the areas of all transitions. A mean profile is obtained by calculating the mean for each transition of all normalized areas across all runs. The mean profile is then normalized. For each peptide in each run, the MPRA is the dot product between the vector containing the normalized areas for each transition and the mean profile calculated previously. This score reflects how similar to each other are the relative peak areas across the entire dataset. The objective of the MPRA is to detect interferences. By optimizing it we can maximize the similarity between the data used to quantify a peptide. By doing this the resulting choice of transitions will be tailored to the dataset to be analyzed, producing the lowest number of interferences in the entire dataset, and providing the solution where the remaining noise has the least impact on the quantification.

For a given peptide using a given set of transitions in a given MS run:

$n$ = total number of transitions in the set
$r$ = total number of MS runs in the dataset
$a_i^j$ = area under the curve between the integration boundaries for transition $i$ in run $j$
$b_i^j$ = normalized area for transition $i$ in run $j$

$$b_i^j = \frac{a_i^j}{\sum_{i=1}^{n} a_i^j}$$

$c_i = mean\ area\ for\ transition\ i\ across\ all\ MS\ runs\ in\ the\ dataset$

$$c_i = \frac{\sum_{j=1}^{r} a_i^j}{r}$$

$c_i' = normalized\ mean\ area\ for\ transition\ i\ across\ all\ MS\ runs\ in\ the\ dataset$

$$c_i' = \frac{c_i}{\sum_{i=1}^{n} c_i}$$

$u^j = vector\ of\ all\ b_i^j\ for\ MS\ run\ j$

$v = vector\ of\ all\ c_i'$

$MPRA^j = MPRA\ score\ for\ MS\ run\ j$

$MPRA^j = u^j \cdot v$

$MPRA = MPRA\ score\ for\ the\ entire\ dataset$

$$MPRA = \frac{\sum_{j=1}^{r} MPRA^j}{r}$$

- Spectral library similarity (SLS) score

For a given peptide and a given subset of transitions, the spectral library similarity is calculated based on the dot product between the intensity of each transition in the spectral library and the areas integrated from the DIA signals. Then the dot product value undergoes a transformation (Supp. Fig. 6B). The aim of this transformation is 1) to set a cut-off value defined by the user under which the SLS score will be low. 2) To examine how similar the measured DIA signals are to the signals present in the spectral library. However, due to instrumental variations over time, changes in instrumental calibration and the use of different acquisition methods (DDA and DIA) the fragmentation patterns observed in the spectral library and the ones obtained when analyzing the samples in DIA might not be exactly the same. In order to be more tolerant to these small changes, another threshold is determined above which the spectral library similarity score will be equal to 1.,e.g. two sets of transitions that have a high dot product will both obtain a spectral library similarity score equal to one. This ensures that both sets of transitions are scored equally instead of biasing one of the two with the dot product that does not take into account the variation of the fragmentation patterns.

- Intensity and intensity product chromatographic-scores

For a given peptide and a given run, the intensity chromatographic score is equal to the sum of the intensities of all transitions at a given time point normalized by the maximum value of the summed intensity across the entire chromatogram. To calculate the intensity product chromatographic score, first 1 is added to the areas of each transition to avoid missing values or values equal to zero. For each time point the areas are multiplied together, log2 transformed, and normalized by the maximum value across the entire chromatogram.

- Potential peak score

In order to define a signal as a potential peptide chromatographic peak, at least 3 transitions should have an intensity higher than the level of noise for at least N number of consecutive points. N is defined by the user and needs to be adapted for each instrument. The level of noise was estimated in the following way: for a given peptide, the level of noise was set as the median of the all the lowest intensity values among all transition at each time point, plus 2 times the standard deviation of these values. This parameter is necessary when analyzing data from Q-TOF instruments as the background noise is higher in these datasets. For Overlap DIA Orbitrap data the noise if very low and the level of noise was set to zero. If the criteria described here is met the Potential peak score is equal to 1, and 0 if not.

Score combination:

Avant-garde is very conservative and uses a novel ensemble-driven scoring strategy. The combined score from Avant-garde is not calculated by adding subscores like other tools. We avoided adding subscores because it can lead to a high number of false positives due to the fact that the combined score can be mainly influenced by one single very good-scoring subscore. Avant-garde changes the way peak groups are scored. The main idea behind Avant-garde is to reduce noise to the minimum, thus obtaining very high-quality signals. We manage this by penalizing peptides having any low-scoring metric. All Avant-garde scores have values between 0 and 1. To combine them each metric is weighted by an exponent and multiplied together. This means that if a peptide does not score well with any given metric the combined score will be severely penalized. Three different combinations are used in AvG for different purposes. The combined scores are called “AvG chromatographic score” for the refinement of peak integration boundaries, “AvG fitness score” for the refinement of the transition selection and “AvG score” for the scoring of peaks.

1. For transition refinement:

For each step (or generation) the genetic algorithm selects a population of randomly chosen subset of transitions, scores each one with a fitness function and selects the best-scoring solutions as the starting point for the next generation.

The fitness function used by the genetic algorithm was defined as follows:
If a randomly selected set of transitions has a number of transitions below a minimal number defined by the user then $Fitness.Score = 0$.
If for a randomly selected set of transitions at least n transitions are not among the top N most intense transitions in the spectral library then $Fitness.Score = 0$. Where n and N are user-defined values.
Otherwise:

*AvG Fitness Score= (1-α-β) x (MPRA + PSS)+ α x IntensityScore + β x MassErrorScore + 0.05xSLS*

Where α and β are user-defined values (default to 0.05).

For the fitness function, more weight is given to the PSS and the MPRA score, given that they are the most sensitive metrics to detect interferences. This enables to confidently identify interferences and remove them in the entire dataset. The intensity, mass errors and library score have a smaller weight on the fitness score. They are used to decide between possible solutions with similar peak shape similarity and MPRA scores. When two solutions are possible the fitness function is designed to choose the one providing the highest intensity, lower mass deviations and matching the best the spectral library.
Often interferences can overshadow the signal from the analyte of interest, especially for low abundant peptides. Reducing the influence of interferences with high intensity on the fitness score by giving a small weight to the intensity score is extremely important. Additionally, the algorithm ensures that at least n transitions are among the N most intense fragments in the spectral library (n and N are user-defined values) guaranteeing that the solution will have intense signals and ensures that the transition selection step will not have a negative impact on the sensitivity of the quantification.

2. For peak boundaries refinement :

In order to refine a peptide identification and its peak boundaries, Avant-garde uses chromatographic scores that are combined to form the *AvG chromatographic score*. The maximum value of the *AvG chromatographic score* corresponds to the peaks' retention time. The boundaries correspond to the retention time where the intensity score is at 4% of the maximum value of the summed intensities of all transitions in the subset.
In order to robustly combine the subscores and estimate the trend of the scores without being too susceptible to rapid changes and isolated anomalies, the scores were first transformed using a moving median. A moving median is a function which replaces each data value with the median of neighboring values. The combined score, termed *AvG chromatographic score*, is calculated in the following way for each time point:

*AvG chromatographic score = moving.average($SLS^3$ x $MPRA^3$) x Intensity.Product.$Score^3$ x moving.average(MassErrorScore) x PotentialPeak*

3. For peak scoring and FDR calculation:

After the transition refinement and peak boundaries refinement steps, each peak group is scored in order to filter the data and control the FDR. The data being scored here are the chromatogram traces between the new integration boundaries. For each peptide in each run, the AvG score is calculated as:

*AvG score = $PSS^{9.5}$ x $SLS^{4.5}$ x $MassErrorScore^{2.5}$ x $MPRA^{0.5}$*

The exponents were found using the HEK293T dataset where one thousand peptides and their corresponding decoys were extracted and the data was curated by AvG. We then determined the set of exponents (multiples of 0.5 between 1 and 10) that increased the separation between target and decoy peptides. These exponents were empirically determined and fixed to this value so we can compare different datasets to each other. The intensity score was not included in the score at this stage in order to avoid penalizing low-abundant peptides and give a strong influence to high-intensity interfered transitions.

<u>AvG score and FDR estimation</u>

In order to define the AvG score that is used to estimate the FDR we used two datasets that were acquired on two different instruments. The first one was acquired on a Q-Exactive HF (Thermo Fisher Scientific) plus instrument. For this dataset, one thousand peptides, and their corresponding shuffled-sequence decoy peptides, from the HEK293T digest data were randomly selected in 15 MS runs (Supp. Figure 4 and 6). The second dataset was acquired on a Q-TOF instrument (Triple-TOF 6600, Sciex). We used a subset of the LFQBench data that contained 4000 target and their corresponding 4000 decoy peptides in 6 runs (Supp. figure 7 and 8).
Avant-garde was used to curate the data and score each peptide in each of the 15 MS runs.
The AvG score was defined as:
AvG score = $PSS^{a1} \times SLS^{a2} \times MassErrorScore^{a3} \times MPRA^{a4}$

An optimization algorithm was used to find the optimal values of the exponents (a1 to a4) of each subscore that enabled to obtain the largest number of validated measurements. This algorithm chose a random set of 4 numbers that corresponded to the exponents in the equation. The random values were a multiple of 0.5 between 1 and 10. The set of exponents that provided the largest number of validated measurement at an FDR of 1% over 1000 iterations were chosen.

The coefficients found to calculate the AvG score were the following:
AvG score = $PSS^{9.5} \times SLS^{4.5} \times MassErrorScore^{2.5} \times MPRA^{0.5}$

The results obtained by this heuristic approach were compared to the results obtained with a standard linear discriminant analysis classifier. To obtain comparable results we used the logarithmic transformation of the equation above:

$$K = a1 \times \log10(PSS) + a2 \times \log10(SLS) + a3 \times \log10(MassErrorScore) + a4 \times \log10(MPRA)$$

The LDA was used to define the set of coefficients (a1 to a4) that enabled to obtain the best separation between targets and decoys. To obtain an equivalent result as the AvG score, the LDA score was then defined as:
LDA score= $10^K$

However, using the LDA score to reach an FDR of 1 % produced a lower number of validated measurements than the one found using the AvG score. This was expected as the AvG score was designed to optimize the number of validated measurements for an FDR below 1%. The LDA is affected by very low scoring decoy signals that are most likely background noise. In order to improve the results of the LDA, we performed a second round of analysis on a subset of the data. We removed all measurements having a LDA score lower than the 25% percentile of the LDA score for decoy peptides. By removing the lower scoring signals, the 2nd round LDA was better at separating target and decoy signals. This is due to the fact that, in the second round, the characteristics of the decoy population is more similar to the target signals as the background noise was removed (Supp. Figure 5). The LDA coefficients were determined using the reduced dataset but the FDR was determined using the complete dataset. In these two datasets, the AvG score shows similar performance as two rounds of linear discriminant analysis.

In this manuscript, we decided to use the AvG score in all the results presented here. This was done to obtain scores that can be compared from dataset to dataset. Additionally, AvG does not produce any output for signals that have a lower number of transitions than the minimal number allowed or that do not have any potential peak. The latter is often the case for decoy peptides. For data acquired in centroid mode, the background noise level is drastically reduced and consequently the number of reported decoys is lower than for data acquired in profile mode. Only signals that have the characteristics of a peptide signal obtain an AvG score. Hence this can produce datasets with a very small population of decoys having an AvG score. In this case it is not possible to perform an LDA classification on these datasets. This does not mean that the decoys were ignored. It means that the quality of the decoy is so low that they do not obtain an AvG score. By fixing the exponents used to calculate the AvG score, we were able to estimate the FDR in all datasets without the need to perform an LDA for each one.

In this manuscript the data acquired on a Q-Exactive series instrument using an overlapped DIA method was filtered using the AvG score and using the following thresholds: SLS >0.7, mass error score>0.7, PSS >0.85, MPRA > 0.9, AvG score >0.1. This guaranteed that all the reported signals had at least a minimal level of signal quality. We then calculated the FDR to verify that it was below 1%. For Q-TOF stepwise DIA data, the data was filtered using the following thresholds: SLS >0.7, mass error score>0.7, PSS >0.85, MPRA > 0.9, AvG score >0.61.
